# Population genomic evidence of adaptive response during the invasion history of *Plasmodium falciparum* in the Americas

**DOI:** 10.1101/2022.10.30.514183

**Authors:** Margaux J. M. Lefebvre, Josquin Daron, Eric Legrand, Michael C. Fontaine, Virginie Rougeron, Franck Prugnolle

## Abstract

*Plasmodium falciparum*, the most virulent agent of human malaria, spread from Africa to all continents following the out-of-Africa human migrations. During the transatlantic slave trade between the 16^th^ and 19^th^ centuries, it was introduced twice independently to the Americas where it adapted to new environmental conditions (new human populations and mosquito species). Here, we analyzed the genome-wide polymorphisms of 2,635 isolates across the current *P. falciparum* distribution range in Africa, Asia, Oceania, and the Americas to investigate its genetic structure, invasion history, and selective pressures associated with its adaptation to the American environment. We confirmed that American populations originated from Africa with at least two independent introductions that led to two genetically distinct clusters, one in the North (Haiti and Columbia) and one in the South (French Guiana and Brazil), and the admixed Peruvian group. Genome scans revealed recent and more ancient signals of positive selection in the American populations. Particularly, we detected positive selection signals in genes involved in interactions with host (human and mosquito) cells and in genes involved in resistance to malaria drugs in both clusters. We found that some genes were under selection in both clusters. Analyses suggested that for five genes, adaptive introgression between clusters or selection on standing variation was at the origin of this repeated evolution. This study provides new genetic evidence on *P. falciparum* colonization history and on its local adaptation in the Americas.

## Introduction

Despite the remarkable advances in medical research and treatments during the 20^th^ century, infectious diseases remain among the leading causes of death in low- and middleincome countries (Geneva: World Health Organization 2022). One of the reasons is the frequent emergence of new infectious diseases and the re-emergence of old ones (Zumla and Hui 2019; Sabin et al. 2020). Emerging infectious diseases are diseases that have recently increased in incidence, in geographic or host range, are newly recognized, or are caused by new pathogens. One key aspect of their emergence is the pathogen adaptability to new environmental conditions (e.g. ecosystems, hosts, treatments, vectors). Explaining how pathogens can adapt to new environments is a prerequisite to better understand the emergence of infectious diseases.

For many pathogen species, the experimental investigation of genetic adaptation to new environments is difficult due to the challenges associated with culturing species in laboratory conditions, especially species with complex life-cycles or with long generation times. An alternative strategy is to analyze well-documented past emergence events that provide a natural “experiment” to identify the genetic basis of genetic adaptation associated with the colonization of novel environments. During the evolution of human populations, new infectious diseases have recurrently emerged. Therefore, there are now several examples of well-documented past emergence events that might serve as models to analyze how pathogens adapted to new environmental conditions (Wolfe et al. 2007; Choi and Thines 2015). *Plasmodium falciparum* colonization of the Americas is one of such examples.

*P. falciparum* is a unicellular eukaryote parasite that causes the most severe form of human malaria, leading to the death of approximately half a million people every year, mainly among <5-year-old children (World Health Organization 2020). Therefore, it is a major public health issue. During its life cycle, *P. falciparum* successively infects two hosts: a vector of the genus *Anopheles* (mosquito) and *Homo sapiens sapiens.*

During its evolutionary history, *P. falciparum* emerged and colonized new geographical areas several times (Tanabe et al. 2010). One of the most recent major events was its introduction to the Americas from Africa during the transatlantic slave trade that lasted from the 16^th^ to the 19^th^ century (Anderson et al. 2000; Yalcindag et al. 2012; Rodrigues et al. 2018). During its colonization of the New World, *P. falciparum* encountered a new human host environment (Amerindian populations, European populations, African and admixed populations) and new vector species (*Anopheles darlingi,* one of the main vector species in South America (Zimmerman 1992; Laporta et al. 2015), and *Anopheles albimanus,* the main vector in Central America (Zimmerman 1992; Frederick et al. 2016)). Compared with *Anopheles gambiae sensu lato* species complex and *Anopheles funestus* (Gillies and Coetzee 1987) (the main African malaria vector species), *An. darlingi* belong to a different subgenus that might have diverged between 80 and 100 million years ago (Moreno et al. 2010; Neafsey et al. 2015; Martinez-Villegas et al. 2019). Moreover, in the last few decades, *P. falciparum* went through another dramatic change in its environment: the use of antimalarial drugs (Wongsrichanalai et al. 2002; Mita et al. 2009) to prevent or cure malarial infections. Some of these drugs are still massively used and exert very strong selective pressures on the parasite. Different populations around the world have developed resistance to these drugs and some new *P. falciparum* genotypes have spread (Mita et al. 2009; Plowe 2009).

The environmental changes during or after the colonization of the Americas by *P. falciparum* likely resulted in powerful selective pressures that forced the parasite to adapt to these new local conditions and to evolve toward different phenotypes and genotypes. Previous studies on few candidate genes identified some genes that may have played a role in the successful *P. falciparum* colonization of the Americas, for instance, the P47 (Molina-Cruz and Barillas-Mury 2014; Canepa et al. 2016; Tagliamonte et al. 2020) and P48/45 genes (van Dijk et al. 2001), two genes involved in the avoidance of the mosquito immune system. Moreover, some studies showed that, in the New World, some *P. falciparum* genes that encode proteins implicated in red blood cell invasion, such as the erythrocyte binding antigen (*eba*) surface ligand are either under stronger selection (Yalcindag et al. 2014) or differentially expressed (Lopez-Perez et al. 2012) compared to the African populations. However, no study has performed a genome-wide analysis of *P. falciparum* response to these new environmental conditions. Yet, due to the large amount of genomic data available, *P. falciparum* offers a unique opportunity to study its genomic adaptation in the context of the colonization of a novel environment. Moreover, the two independent *P. falciparum* introductions in the New World from Africa during the transatlantic slave trade (Yalcindag et al. 2012) may be regarded as two potentially independent replicates of the same “natural experiment”. It has been suggested that the first introduction occurred in the south of the continent (through Brazil) and the second, more recent, in the north (through Colombia). These two independent introductions likely coincided with the two main slave trade routes from Africa to the Americas: one organized by the Portuguese empire to bring slaves to the current Brazil (600 years ago), and the other organized by the Spanish empire to Colombia (300 years ago) (Yalcindag et al. 2012). These two independent introductions can give information on whether evolution followed the same path (repeated or parallel evolution) and whether the same genes responded to the novel environmental conditions. As the populations that resulted from these two introductions have historically exchanged migrants (Yalcindag et al. 2012), it is also possible to determine the importance of migration in the adaptive evolution of the introduced populations.

In the present study, we compiled four whole genome polymorphism datasets of *P. falciparum* from the Americas, the source area (Africa), and other world areas (Asia and Oceania). Using these data, we studied the population genetic structure and colonization history of the American *P. falciparum* populations. We then scanned the genomes to identify genes and genomic regions that may have responded to the selection imposed by the colonization of novel environments in the New World. As selection may have occurred at different time points in the history of the introduced populations, we used complementary methods to identify recent and also more ancient signals of selection. Then, for genes that showed signals of selection in distinct American populations, we determined whether alleles were independently selected within each location or whether they resulted from a process of adaptive migration between populations.

## Results

### Compilation of whole genome polymorphism data

We obtained whole-genome polymorphism data for *P. falciparum* from publicly available datasets. Specifically, we retrieved the single nucleotide polymorphism (SNP) data (VCF files) from the *MalariaGen Project* (Pearson et al. 2019) that includes 7,113 isolates from 29 countries, particularly Africa (3,877 isolates), Asia (2,942 isolates), Oceania (231 isolates), and South America (39 isolates), and also laboratory samples (16 isolates) and samples collected from travelers (8 isolates). As this dataset only included two countries from the Americas (Colombia and Peru), we also used population data from Brazil (23 isolates) (Moser et al. 2020), French Guiana (36 isolates) (Pelleau et al. 2015), and Haiti (21 isolates) (Tagliamonte et al. 2020). The total dataset before filtering included 7,193 strains from 32 countries in the Americas, Africa, Asia, and Oceania.

For the *MalariaGen* dataset, we quality filtered the SNP data and samples following the recommendation by Pearson et al. (2019). We kept only high-quality bi-allelic SNPs from the core genome (*i.e.,* regions present in all individuals, as defined by Pearson et al. (2019)) and 5,970 samples that corresponded to the analysis set as defined by Pearson et al. (2019). We excluded the only sample from Mozambique, from a traveler, to avoid bias due to a population made of only one individual. For all populations, we removed samples with >20% missing data (11 samples) and with multi-strain infections (*i.e.* multiple *P. falciparum* genotypes), reported as F_WS_ >0.95 (*Supplementary Fig. S1*). In addition, for all pairs of strains displaying excessive relatedness (pairwise-IBD>0.5), we excluded one strain (*Supplementary Fig. S1*). The final sample included 2,635 individuals (*Fig. 1A, Supplementary Table S1, Supplementary Fig. S2*).

**Figure 1:**
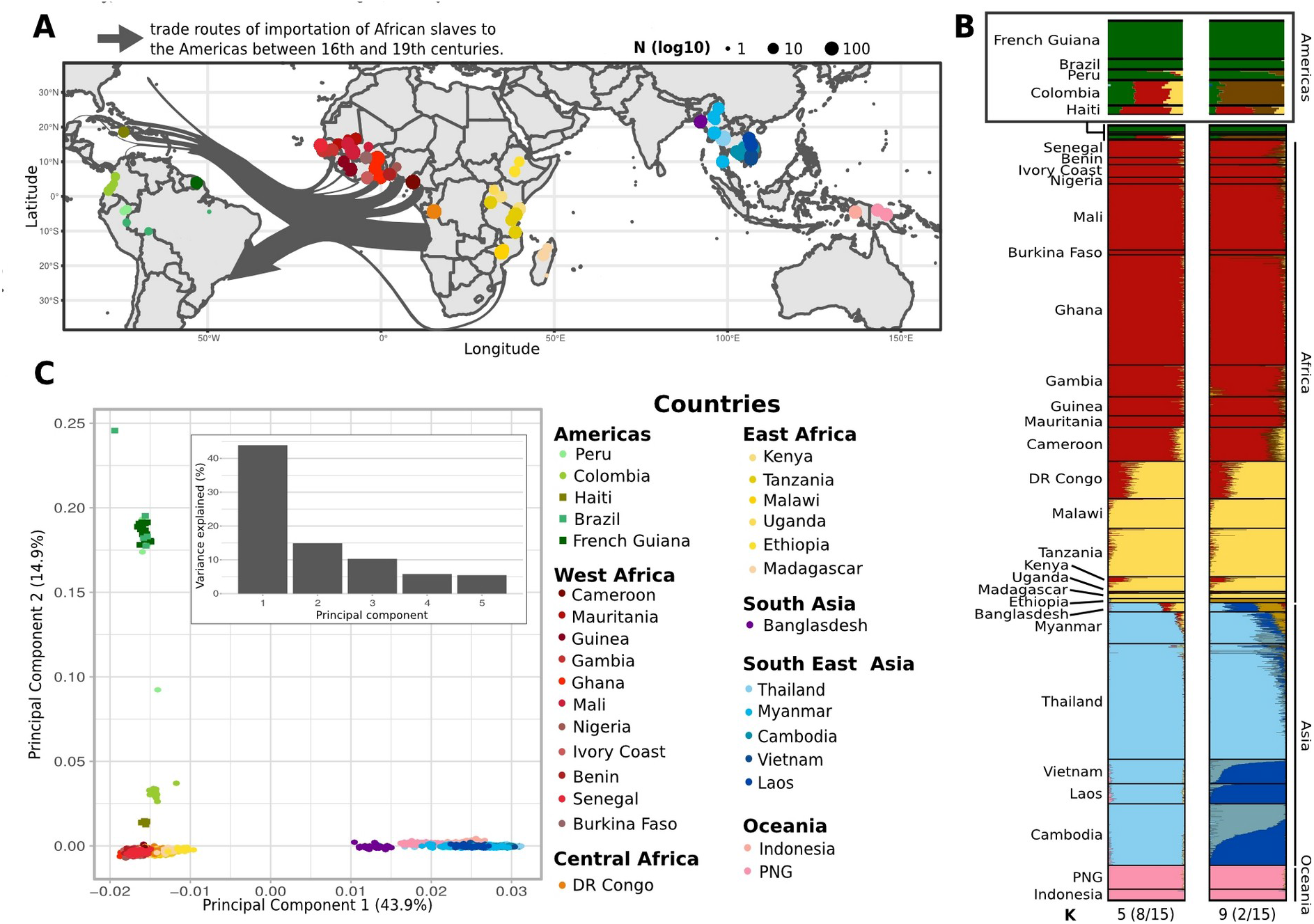
Geographical origin and genetic structure of *Plasmodium falciparum* isolates. **A.** Geographical origin of the 2,635 *P. falciparum* isolates per region: South America (n=48), West Africa (n=1,046), Central Africa (n=138), East Africa (n=348), South Asia (n=36), South East Asia (n=885), and Oceania (n=112). The circle size is proportional to the sampling effort per site (log10 N). Arrows indicate the slave trade waves, and their width is proportional to the number of slaves transported (source: www.slavevoyages.org, accessed on 25/10/2021). **B.** ADMIXTURE results for K=5 and K=9 clusters (indicated at the bottom, with the clustering convergence rate for 15 replicated runs). DR Congo: Democratic Republic of Congo. PNG: Papua New Guinea. **C.** Graphical representation of the Principal Component Analysis results. Each color represents a country; circles correspond to *P. falciparum* genomes obtained from the MalariaGen Project (Pearson et al., 2019); squares represent the newly added data for this study (Haiti, Brazil, and French Guiana). The histogram represents the percentage of the genetic variance explained by the five first principal components.

Then, from this dataset, we removed SNPs with >10% of missing data. We set a minimum allele frequency (MAF) filter at 0.01%, to delete polymorphisms associated with sequencing errors, and a minimum coverage depth at 15X. In the end, our dataset included 78,036 SNPs for 2,635 samples, with a mean SNP density of 3.69 per kilobase. The filtering steps are described in *Supplementary Figure S2.* Following Pearson et al. (2019), we defined eight geographic regions for the analyses: the Americas (SAM), West Africa (WAF), Central Africa (CAF), East Africa (EAF), South Asia (SAS), West Southeast Asia (WSEA), East Southeast Asia (ESEA) and Oceania (OCE).

### Two distinct admixed genetic clusters of *P. falciparum* in the Americas

To obtain insights into the genetic relationships among isolates from all over the world, we first investigated *P. falciparum* population structure using two complementary approaches: *i*) a genetic ancestry analysis with the ADMIXTURE v1.3.0 software and *ii*) principal component analysis (PCA). For these analyses, we used a dataset of 41,921 SNPs from 2,434 individuals after linkage disequilibrium (LD) pruning, MAF filtering, and missing data filtering (see Material and Methods for more details).

The genetic ancestry analysis with ADMIXTURE identified nine distinct genetic pools as the optimal number of clusters (K) of *P. falciparum* populations worldwide (*Supplementary Fig. S3*). With K=2, Asian populations formed one cluster, and African and American populations the other (*Supplementary Fig. S4*). From K=5 onward, the American populations became distinct from the others (*Fig. 1B*). With K=9, isolates from Brazil and French Guiana formed a distinct genetic cluster (dark green in *Fig. 1B),* while isolates from Peru, Colombia, and Haiti displayed evidence of admixed ancestry, especially with African populations (brown in *Fig. 1B*). We obtained similar results by PCA (*Fig. 1C*). The first principal component (PC), which represented 43.9% of the variance explained, separated the Asian populations from the African and American populations. The second PC (14.9% of variance explained) split African isolates from American isolates, and also showed the intracontinental structuring of American isolates. Indeed, Brazil and French Guiana isolates were the most distant from African isolates, and we could not identify any finer subdivision. Conversely, isolates from Colombia and Haiti were quite distinct and very close to Africa. Peruvian isolates were in the middle between the cluster formed by Haitian and Colombian (SAM North cluster) isolates and the cluster formed by Brazilian and French Guianan (SAM South cluster) isolates.

To further investigate the genetic relationships between the American populations and populations from the rest of the world as well as their colonization history, we estimated population trees and networks using two approaches implemented in TreeMix (Pickrell and Pritchard, 2012) and ADMIXTOOLS2 (Maier et al., 2022), respectively. These two methods use the shared and private genetic ancestry components to infer the population branching, while taking into account also historical migration and admixture events between populations. For both analyses, we used genome data of *Plasmodium praefalciparum*, the closest known relative of *P. falciparum,* as an outgroup to root the tree/network, and a LD-pruned dataset composed of 20,943 SNPs for 2,638 individuals.

The TreeMix analysis identified three migration edges as optimal for our dataset (*Supplementary Fig. 5A-B*). The resulting TreeMix consensus tree (*Fig. 2A*) confirmed the global genetic structuration given by the PCA and ADMIXTURE analyses, particularly the close relationship between the African and American populations. Many African populations were very closely related, with very short branches. In other words, they were all part of the same cluster that did not drift much in recent times. The tree showed that the two American clusters (SAM North and SAM South) connected to different positions in the tree. SAM North populations (Haiti - Colombia) were more closely related to populations of West Africa, whereas SAM South populations (Brazil - French Guiana) and Peru were branched deeper in the tree as a sister group to the African cluster. Moreover, the American populations were characterized by a strong drift effect, marked by longer branches. This reflects founder effects. Our results for the American populations also indicated 25% of interbreeding with an older or unsampled population (on the *P. praefalciparum* branch), and more recent mixing (up to 36.35%) between the Colombian branch and the cluster composed of Peruvian, Brazilian and French Guianan populations (*Fig. 2A*).

**Figure 2:**
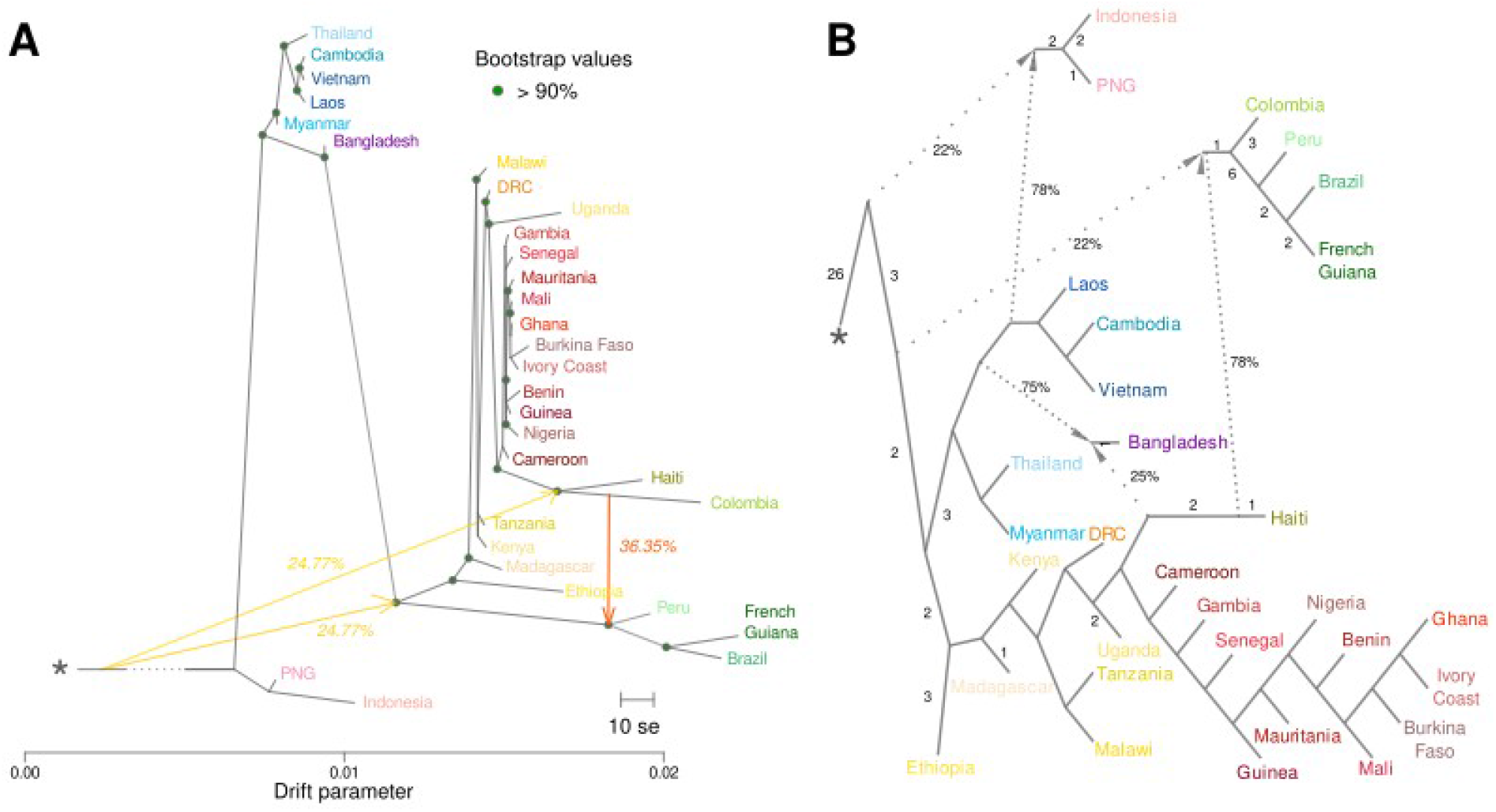
Relationships and admixture proportions between *P. falciparum* populations. **A.** TreeMix tree of *P. falciparum* populations with three migration edges (arrows), rooted with *P. praefalciparum*. Filled circles represent the bootstrap values supported by more than 90%. The scale bar shows ten times the mean standard error (se) of the entries in the sample covariance matrix. Arrows show the migration edges between tree branches. The migration weight is indicated by the yellow (low) to red (high) color gradient and the estimated value is shown as a percentage on the migration edge. **B.** Admixture graph of *P. falciparum* populations, with three admixture events, rooted with *P. praefalciparum*. Admixture events (and the estimated percentages) are shown with dash arrows that connect the admixed population with the two source population branches. The number on each branch represents the branch length that indicates the amount of drift accumulated along that branch (in *f*-statistic units, multiplied by 1,000 and rounded to the nearest integer). Branches without a number are branches with a drift score = 0. PNG: Papua New Guinea, DRC: Democratic Republic of Congo. The asterisk represents the *P. praefalciparum* outgroup.

The ADMIXTOOLS2 analysis also identified an optimal number of admixture events equivalent to the one found with TreeMix (n=3). This network of *P. falciparum* populations confirmed the results of the population structure analyses (*Fig. 2B*). Haiti and the common ancestor of Colombia, Peru and SAM South cluster shared 78% of ancestry, although 22% came from an older or unsampled population. Moreover, the Haitian population seemed to be the result of important interbreeding between African and South American populations, as suggested by ADMIXTOOLS2 (*Fig. 2B*) and also ADMIXTURE with K=9 (*Fig. 1B*).

Altogether, these results suggest that the American populations are subdivided into two distinct genetic clusters: a southern cluster that includes Brazil and French Guiana (SAM South) and a northern cluster that includes Colombia and Haiti (SAM North). The TreeMix and ADMIXTOOLS2 analyses also suggested that all southern populations (SAM South cluster and Peru) are the result of an ancestral large admixture event between the northern and southern clusters and also with unsampled populations.

### Founder effect associated with *P. falciparum* colonization of the Americas

To assess the demographic history of the American populations, and particularly whether they went through bottleneck events during the colonization process, we calculated the Tajima’s *D* values (Tajima 1989), and analyzed the effective population size (*N_e_*) changes through time using *Stairway Plot 2* (Liu and Fu 2020). For these analyses, we only kept the American populations grouped by cluster (excluding the highly admixed Peruvian population), and three African populations (Democratic Republic of Congo, Tanzania, and Senegal) and one Asian population (Myanmar) for comparison. The polarized dataset included 31,892 SNPs for 500 samples, from which we could generate an unfolded site frequency spectrum (SFS).

The Tajima’s *D* values of the American populations were positive and not different between the SAM genetic clusters (Wilcox test, Bonneferroni adjusted *p*-value: 1.32e-01). Conversely, the African and Asian populations displayed Tajima’s *D* values close to zero (*Fig. 3A*). This relative lack of rare alleles or excess of shared variants in the American populations, characterized by positive Tajima’s *D* values, suggests a historical demographic contraction. Analysis of the population size changes, based on the SFS (*Fig. 3B),* indicated that all American populations showed similar patterns, characterized by a decline or a bottleneck in the population that started between 3,000 and 3,600 generations ago. By considering six generations per year for *P. falciparum*, this corresponded approximately to 600 and 500 years ago (*i.e.* the time of slaves’ arrival in the New World). More recently, SAM North (Haiti - Colombia) underwent a decline (600 generations ago, ~ 100 years ago), followed by a bottleneck (240 generations ago, ~ 40 years ago). SAM South (Brazil - French Guiana) populations declined 300 generations ago (~ 50 years ago). In comparison, the African and Asian populations did not show any decline in the same period (*Fig. 3B*).

**Figure 3:**
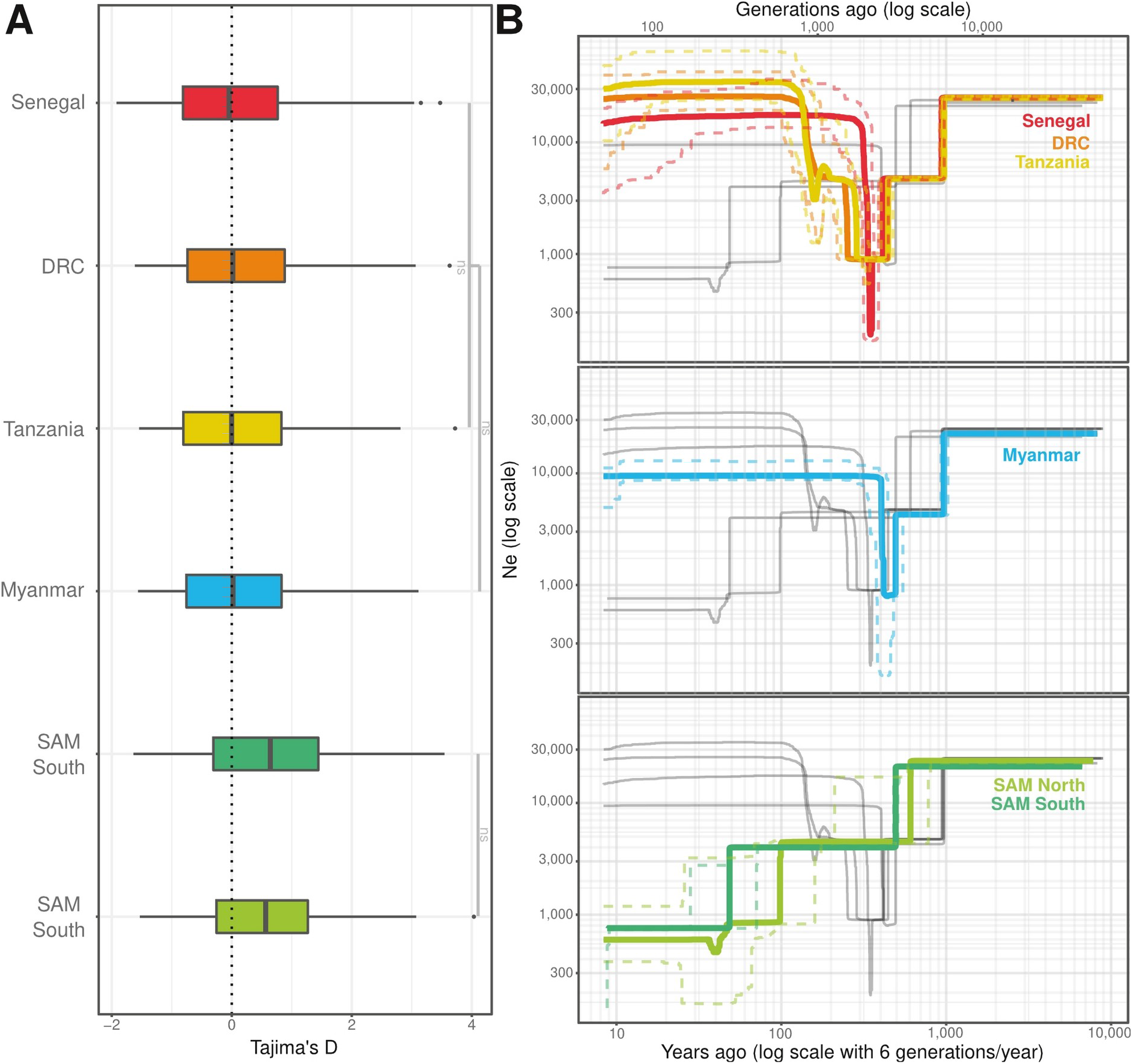
Demographic history of *P. falciparum* in South America, compared with African and Asian populations. **A.** Comparison of the Tajima’s D values of the two American genetic clusters, three African populations (Senegal, Tanzania, and DRC), and one Asian country (Myanmar). **B.** Effective population size inferred from *Stairway* plot 2 for the two American clusters, three African populations (Senegal, Tanzania, and DRC), and one Asian country (Myanmar). The solid lines correspond to the median and the dashed lines represent the 95% confidence interval. Gray lines represents the other populations. Note that the axes are in the log_10_ scale. DRC: Democratic Republic of Congo.

### Genomic evidence of recent and ancient local adaptations in the American populations

We first used haplotype-based tests (*XP-EHH* and *Rsb*) to detect evidences of recent positive selection in the *P. falciparum* genomes in the Americas compared with Africa. *XP-EHH* and *Rsb* allow detecting signals of selection that are specific to the introduced populations compared with the source (here, the African populations). As these tests are based on haplotype length and linkage disequilibrium, they tend to detect recent or ongoing positive selection events because signals of more ancient selective events would have been broken by recombination over time (Voight et al. 2006; Sabeti et al. 2007; Tang et al. 2007). We performed all tests for the SAM North and SAM South clusters independently, and used Senegal as the reference population from the native African zone. The dataset for these analyses included 78,036 SNPs for 98 samples.

In the SAM North cluster (Colombia - Haiti), the *XP-EHH* and *Rsb* tools detected respectively 20 and 19 genes with significant SNPs in their coding sequence (CDS) or untranslated region (UTR) (*Supplementary Fig. S6,* and *Fig. 4),* of which 17 were in common. In the SAM South cluster (Brazil - French Guiana), 16 genes displayed significant SNPs in the CDS or UTR regions with *XP-EHH (Supplementary Fig. S6*) and 16 with *Rsb* (*Fig. 4);* 14 genes were in common. All candidate genes identified as potentially under positive selection in the American clusters are listed in *Supplementary Tables S2-4.*

**Figure 4:**
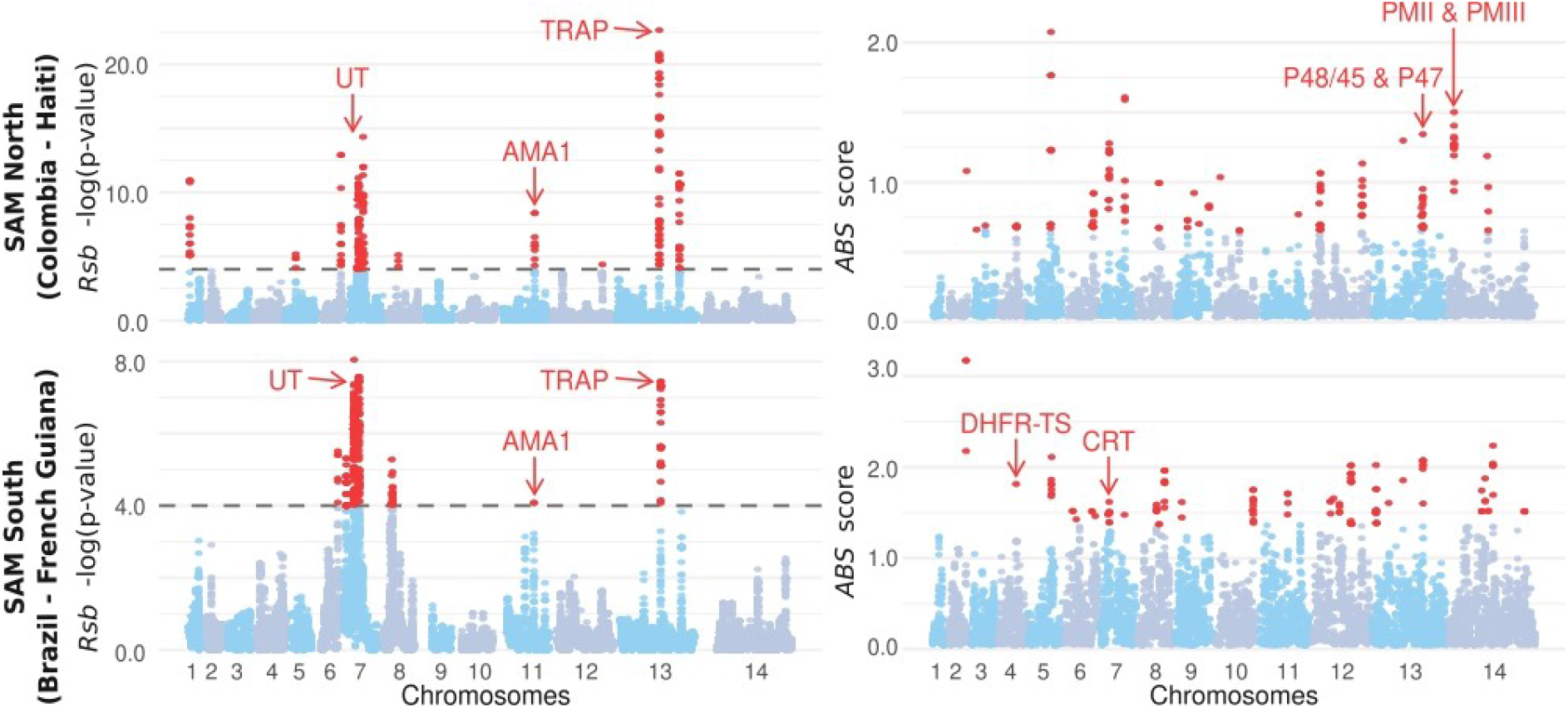
Evidence of selective sweeps in the two American genetic clusters. Manhattan plots showing the *Rsb* and ABS selection scans for the SAM North (Colombia - Haiti) and SAM South (Brazil - French Guiana) clusters. For the *Rsb* scores, the dotted lines represent the threshold significance value -log(p-value) = 4. Red, SNPs marking a selective sweep in the SAM cluster (negative *Rsb* values). For the *ABS* scores, all values in red represent the top 1% of values, evidence of past positive selection events. UT: ubiquitin-protein transferase, DHFR-TS: dihydrofolate reductase-thymidylate synthase, CRT: chloroquine resistance transporter, AMA1: apical membrane antigen-1, TRAP: thrombospondin-related adhesive protein, PMII: plasmepsin II, PMIII: plasmepsin III.

These selection tests provided primary evidence of recent selection signals. However, it is likely that *P. falciparum* colonization of the American continent started more than 500 years ago (~3,000 generations ago, depending on the estimated date of its introduction and of its expected generation time) (Yalcindag et al. 2012). During this time, the parasite might have responded to the selective pressures during the first generations of its colonization history and these “ancient” signals of selection may have been hidden by recombination events occurring at each generation, breaking up linkage disequilibrium tracks along the genome. Therefore, to detect more ancient positive selection signals, we used the ancestral branch statistic (*ABS*), an *FST*-like statistic in which two closely-related populations from the same American cluster were compared with two outgroup populations (one from Senegal, Africa, and one from Myanmar, Asia) in a quartet population system (Cheng et al. 2017). This approach allowed detecting signals of selection in the ancestral branch that linked the American populations of each cluster to the African/Asian populations. The outgroup (Senegal and Myanmar) populations were the populations with the least amount of missing data in the two continents. We used the dataset already exploited for the *XP-EHH* and *Rsb* analyses, to which we added 103 isolates from Myanmar.

We found significant signals of positive selection for 109 and 188 genes with CDS or UTR regions included in outlier windows that putatively underwent positive selection in SAM North (*Supplementary Table S2 and S4*) and in SAM South (*Supplementary Tables S3 and S4*), respectively. We identified several genes implicated in interactions with the hosts (*Anopheles* spp. and/or humans) and with drug resistance (*Fig. 4*). None of these genes overlapped with the results obtained with the haplotype-based tests (*XP-EHH* and *Rsb*).

### Cluster-specific selection, parallel evolution, or adaptive migration between the northern and southern American clusters

The two independent waves of introduction in the Americas offered the opportunity to determine whether evolution proceeded similarly in the two clusters to adapt to the new environments. To this aim, we looked for different categories of genes or genomic regions that showed (i) a signal of selection in one but not in the other cluster (cluster-specific selection); and (ii) evidence of positive selection in both clusters.

Most loci (n=80/132 in SAM North and n=154/206 in SAM South) showed evidence of selection in one cluster, but not in the other. For instance, *ABS* showed that some genes involved in the parasite immune evasion from the mosquito immune system (*i.e.* P48/45 and P47) were under selection exclusively in the SAM North cluster. Moreover, in SAM North populations, genes involved in resistance to treatments (plasmepsin II and III, PMII and PMIII) displayed evidence of ancient positive selection events, independently of the SAM South cluster. Lastly, the chloroquine resistance transporter (CRT) gene, which is implicated in *P. falciparum* resistance to chloroquine, was under positive selection only in the SAM South cluster.

Other genes (n=52) showed evidence of positive selection in both clusters: 37 genes with *ABS,* and 14 genes with the other haplotype-based tests (*Supplementary Table S6*). Some of these genes encode proteins implicated in the parasite-host interactions, for instance apical membrane antigen 1 (AMA1) and thrombospondin-related adhesive protein (TRAP), and others in drug resistance (e.g. ubiquitin-protein transferase, UT). The finding that the same genes were under selection in both clusters could be explained by different evolutionary mechanisms: (1) parallel evolution, if the two genes independently responded to similar selective pressures in the two clusters; (2) adaptive migration, if the advantageous allele was first positively selected in one cluster and then migrated to the other cluster where it was also positively selected locally; (3) selection on standing variation, if the same haplotype, from the ancestral population, was selected in both clusters; and (4) selection in the ancestral population, before the cluster divergence (for the genome regions that resulted from introgression or admixture between clusters). To disentangle these different possibilities, for each gene, we used the *Relative Node Depth (RND*) to measure the haplotype similarity around the selected loci (Feder et al. 2005). This statistic takes into account local diversity variations along the genome. A low *RND* value compared with the rest of the genome suggests adaptive migration, while no difference would indicate parallel evolution, selection on the same standing haplotype, or ancestral selection. Most regions fell into the second category. Only five genomic regions had low *RND* values, suggesting adaptive migration between clusters (*Fig. 5A*). The signal was particularly strong for TRAP, a gene involved in the parasite interaction with mosquitoes and humans. Indeed, we found only few, closely related haplotypes in the two American clusters, compared with the high haplotypic diversity in the African populations (*Fig. 5B*).

**Figure 5:**
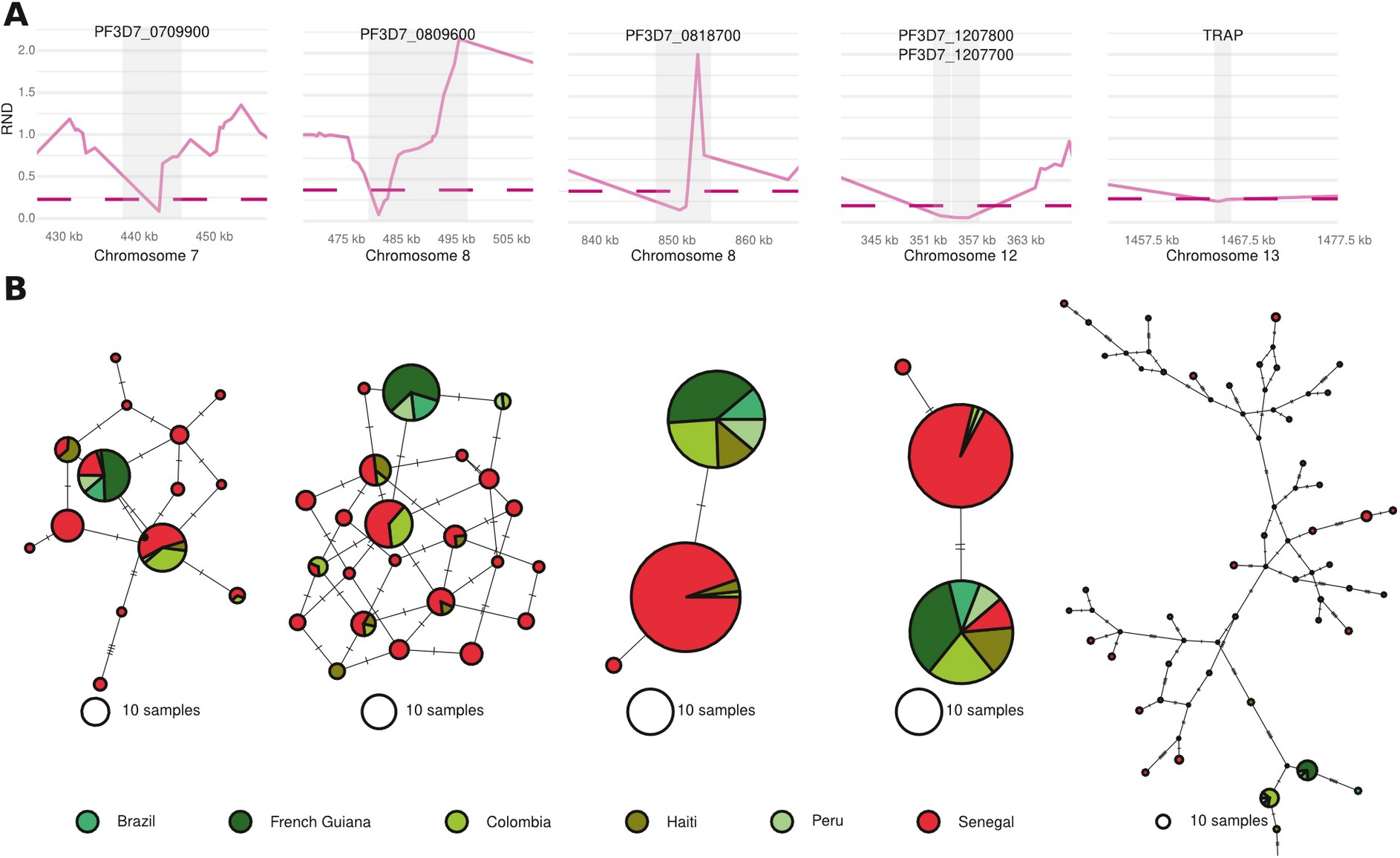
Candidate genes for convergent evolution or introgression between American clusters. **A.** Relative Node Depth (RND) values between SAM North (Haiti - Colombia) and SAM South (Brazil - French Guiana) for five regions. The region in gray represents the gene (gene name on top). The dotted line marks the 5% threshold of the lowest values on the chromosome. **B.** Median-Joining haplotype networks of the regions with the lowest RND values between American clusters. TRAP: thrombospondin-related adhesive protein.

## Discussion

*P. falciparum* was introduced into the Americas from Africa during the transatlantic slave trade from the 16^th^ to the 19^th^ century (Anderson et al. 2000; Yalcindag et al. 2012; Rodrigues et al. 2018). This history of colonization of a new continent offers the opportunity to analyze how this parasite genetically adapted to new environmental conditions (new human host populations with distinct characteristics from the source populations as well as to new mosquito host species and new abiotic conditions). The different waves of introduction that likely occurred during this colonization history can be potentially considered as replicates of the same “natural experiment” to explore the repeatability of adaptive evolution.

### *P. falciparum* population structure and demographic history in the Americas

Genomic information on *P. falciparum* in the Americas allowed us to (re-)explore its population structure and colonization history on this continent. First, we confirmed that the American populations of *P. falciparum* originated from Africa (Anderson et al. 2000; Yalcindag et al. 2012; Rodrigues et al. 2018). Indeed, when we modeled two genetic clusters (K=2) in the ADMIXTURE genetic ancestry analysis (*Supplementary Fig. S3),* we observed a distinction mainly between Asian and African/American populations. American populations split from African populations only with K values >5. This conclusion was confirmed by the PCA (*Fig. 1B*). The population branching obtained with TreeMix and ADMIXTOOLS2 (*Fig. 2*) again indicated that American populations were more closely related to African populations or to *P. praefalciparum*, an African gorilla parasite. Unlike Rodrigues *et al.* (2018), we did not observe any evidence of introgressions from the Asian strains into the American populations. This difference could be explained by the fact that Rodrigues et al. (2018) used mitochondrial markers, whereas we used nuclear SNPs. Indeed, mitochondrial genomes are haploid and clonally transmitted without any recombination among lineages (Galtier et al. 2009; Preston et al. 2014). Therefore, mitochondrial genomes form a single locus potentially subject to selective processes and with very limited resolution and representativeness of the population genetic structure and demographic histories. As these mitochondrial markers have been used to infer ancient colonization or migration events that are undetectable in the nuclear genome (Diez Benavente et al. 2020), these Asian introgressions may reflect ancient gene flow or incomplete lineage sorting.

Our results also confirmed previous observations that American *P. falciparum* populations are subdivided into at least two distinct genetic clusters: one in the North (Colombia and Haiti) and one in the South (Brazil and French Guiana). The Peruvian population was admixed between these clusters. The PCA results (*Fig. 1B*) indicated that the SAM South cluster was more differentiated from the native African populations than the SAM North cluster. Similarly, the ADMIXTURE results showed that the Brazil and French Guiana populations formed a cluster without any evidence of admixture with African populations, even at low K values (*Fig. 1C*). This suggests that the Haitian and Colombian populations arrived in the Americas more recently than the other populations, or have recently been admixed with African populations.

Using 12 microsatellites markers and 384 SNPs, Yalcindag et al. (2012) also found a genetic demarcation between Colombian populations and Brazilian/French Guiana populations, and populations with interbreeding profiles in Peru and Venezuela. Scenario-testing using Approximate Bayesian Computation suggested that this structuring was the result of several independent introductions of *P. falciparum* in South America (Yalcindag et al. 2012). In agreement, our TreeMix and ADMIXTOOLS2 analyses indicated at least two independent introductions into the Americas. However, unlike Yalcindag et al. (2012), our analyses suggested that either both Haitian and Columbian populations were introduced independently of the Peruvian, Brazilian and French Guianan populations, or only the Haitian population (*Fig. 2*). Both analyses also suggested introgression from the SAM North cluster (36.35% from Colombia with TreeMix, and 76% from Haiti with ADMIXTOOLS2) to the south American common ancestor (*Fig. 2*).

Concerning the demographic history of the American populations, our analyses (Tajima’s *D* and *Stairway Plot 2*) suggested that they went through several declines and bottlenecks during or after the colonization of the Americas (*Fig. 3B*). Their consequence is also visible in the longer branch lengths of the TreeMix tree for the American populations compared with populations of the other continents. These results suggest higher genetic drift in American populations compared with the other populations because of founder effects or bottlenecks during the colonization of the new continent and/or intense selection pressure due to new environmental conditions. Indeed, some decline events found with *Stairway plot 2* occurred at times that may correspond to the introduction dates of *P. falciparum* in South America, as proposed by Yalcindag et al. (2012). Conversely, other events were too recent to have been caused by a founding effect following the parasite introduction into the New World (*Figures 3.B*). Specifically, the SAM South (Brazil - French Guiana) cluster experienced an effective population size decrease long after its likely introduction into South America. This recent demographic decline (~50 years ago) might be explained by the selection imposed by antimalarial drugs.

### Evidence of adaptation in the Americas

During the colonization of the Americas*, P. falciparum* faced new environmental conditions that could have exerted strong selection pressures. These adaptation processes have left an imprint in *P. falciparum* genome.

#### Adaptation to new hosts (mosquitoes and humans)

Two genes of the 6-cysteine family (P47 and P48/45), expressed during the stages when *P. falciparum* is present in mosquitoes, showed extreme *ABS* values (*Supplementary Tables S2-3),* indicating positive selection early in the colonization history of the Americas. In *P. falciparum*, P47 allows escaping the vector immune system (Molina-Cruz and Barillas-Mury 2014; Canepa et al. 2016), whereas the proteins encoded by P48/45 play an essential role in reproduction, which takes place in the mosquito (van Dijk et al. 2001). This divergent selection for P47 and P48/45 was previously described in worldwide studies on the polymorphism of these genes (P47: Anthony et al. 2007, and P48/45: Conway et al. 2001). Here, P47 and P48/45 showed evidence of positive selection only in the SAM North cluster (Colombia - Haiti), as already reported by Tagliamonte et al. (2020). In these regions, *P. falciparum* is transmitted by different mosquito species than in Southern South America (Gutiérrez et al. 2008; Boncy et al. 2015), and this could explain the observed between-cluster differences. However, a lack of power to detect selection in the SAM South cluster cannot be fully ruled out.

For some of the genes under selection, the origin of the selective pressure was less evident because they are expressed at different stages of the parasite life cycle, both in humans and mosquitoes. For instance, TRAP allows crossing the cell barriers in both hosts (Akhouri et al. 2004), and AMA1 is important for the invasion of erythrocytes (Triglia et al. 2000) and hepatocytes (Yang et al. 2017). An adaptation of AMA1 to mosquitoes cannot be entirely ruled out because it was recently shown that this protein is involved in the invasion of the mosquito salivary glands (Fernandes et al. 2022).

Other genes (n=127), such as RF1, HAD2 and C3AP3, also may have played a role in *P. falciparum* adaptation to the mosquito and/or to human host, but their function is unknown. Functional analyses may help to better understand *P. falciparum* adaptive processes in the Americas.

#### Adaptation to anti-malarial treatments

Another major selection pressure exerted on *P. falciparum* population in the Americas is related to the use of antimalaria drugs by human populations to prevent infection. *P. falciparum* genome in the Americas includes some evidence of selection concerning known resistance genes (Wongsrichanalai et al. 2002; Mita et al. 2009). We found a common signal in the SAM North and South clusters for UT, a gene potentially involved in resistance to quinine (Sanchez et al. 2014). However, the two American clusters did not present similar profiles for other drug response genes (*Supplementary Tables S2-3*). In SAM North (Haiti - Colombia), we identified selection signals for PMII and PMIII, two genes involved in resistance to the dihydroartemisinin-piperaquine combination (Mukherjee et al. 2018) that currently has been detected only in Asia (Amato et al. 2017; Witkowski et al. 2017). In the SAM South cluster (Brazil - French Guiana), as expected, we found selection signals for the dihydrofolate reductase (DHFR-TS) and CRT genes. DHFR-TS is a gene implicated in the resistance to the sulfadoxine-pyrimethamine (SP) combination (Happi et al. 2005). This treatment was introduced in Venezuela in the 1950s (Gabaldon and Guerrero 1959), and in the 1970s, it was used in various American countries as an alternative to chloroquine, although pyrimethamine-resistant strains had been already detected in some regions (Maberti 1960; Walker and Lopez-Antunano 1968). DHFR-mutant parasites, resistant to SP, indigenously evolved in South America (Mita et al. 2009). The first cases of resistance were reported in Venezuela in 1977 (Godoy et al. 1977) and then in Colombia in 1981 (Espinal et al. 1985). From each of these countries, a distinct resistant lineage spread to South America. Currently, the Colombian lineage is also found in Peru and Bolivia, while the Venezuelan lineage has spread mainly to Brazil and to Bolivia (Mita et al. 2009). The absence of any trace of selection in the SAM North cluster might be explained by the mixing of Colombian and Haitian populations or by the small sample size, which may have reduced the detection power. No SP-resistance haplotype has been observed in Haiti (Carter et al. 2012; Rogier et al. 2020). CRT confers resistance to chloroquine and resistant strains appeared in Colombia and Venezuela in 1960 (Moore and Lanier 1961), independently of other world regions (Mita et al. 2009; Plowe 2009). Then, resistance spread throughout the American continent between the 1960s and the 1980s, with two distinct main genotypes (Mita et al. 2009; Plowe 2009). We did not find any evidence of selection signals for CRT in the SAM North cluster. This was expected because it is thought that the *P. falciparum* Haitian population does not include any chloroquine-resistant strain (Neuberger et al. 2012; Vincent et al. 2018; Rogier et al. 2020), although this remains controversial (Londono et al. 2009; Gharbi et al. 2012). Furthermore, French Guianan populations have recently evolved a compensatory mutation on the CRT gene that makes it sensitive to chloroquine. The frequency of this mutation has increased from 2.7% in 2002 to 58% in 2012 (Pelleau et al. 2015). Such rapid allele frequency increase could suggest a recent adaptation signal that could have been detected with *XP-EHH* and/or *Rsb*. The absence of its detection could be explained by the fact that for these analyses, we mixed the American populations by cluster due to the small sample sizes.

Surprisingly, the majority of significant signals for drug resistance genes were found with *ABS* (a test dedicated to detect ancient signals of selection) and not with *XP-EHH* or *Rsb* (that are supposed to detect more recent signals of selection), despite the fact that the use of antimalarial drugs is recent (Wongsrichanalai et al. 2002; Mita et al. 2009; Plowe 2009). The explanation for that is likely associated to the construction of the statistics. *XP-EHH* and *Rsb* compare the size of extended haplotype homozygosity (EHH) for each SNP between populations (Voight et al. 2006; Sabeti et al. 2007; Tang et al. 2007). As the African and American populations experienced the same drug pressure at about the same time (Wongsrichanalai et al. 2002; Mita et al. 2009; Plowe 2009), they both have long EHH. Therefore, selective sweeps cannot be detected with *XP-EHH* or *Rsb*. Conversely, when using *FST*-based metrics (e.g. *ABS),* haplotype differences between populations are detectable. Indeed, drug resistance-conferring haplotypes are different in the Americas and in Africa and Asia, and have spread to the south American continent (Mita et al. 2009; Plowe 2009), thus increasing the distance with the outgroups (Asia and Africa). This explain why traces of selection on these genes can be detected with *ABS*, although they are not ancient selection events.

#### Cluster-specific selection, parallel evolution, and adaptive migration

Although most genes (n=234) underwent cluster-specific selection, some (n=51) experienced selection in both clusters. It is thought that repeated and parallel evolution is infrequent in populations (Bailey et al. 2015), particularly because there are too many different phenotype combinations and even more genotype combinations that can generate higher fitness in a new environment. Consequently, the probability of parallel evolution for a particular phenotype is considered very low because it is very unlikely that the same combinations of *de novo* mutations might occur twice by chance (Bailey et al. 2015). However, it is more and more acknowledged that much adaptation, especially in the context of a rapid environmental change (e.g. during introduction to a new area), proceeds from the sorting of ancestral standing variation and does not rely completely on *de novo* mutations (Thompson et al. 2019). In the *P. falciparum* populations in the Americas, this might have occurred for most genes showing evidence of selection in the New World, at least for the genome regions that came from two distinct introduction waves in the North and the South (between 20 and 60% of the genome for the SAM South cluster, depending on the analysis). For the other genome regions that originated from admixture or gene flow from the SAM North cluster, selection might have taken place on haplotypes that were already separated by genetic drift.

Convergent gene evolution can occur in independently introduced populations also through adaptive introgression/migration (*i.e.* by introducing the identical adaptive alleles from one population into others where they are selected) (Zhang et al. 2021) or through selection on standing genetic variation (Lee and Coop 2019). In the first scenario, gene flow plays an important role in moving the same allelic variants that share a single mutational origin among populations (Zhang et al. 2021). In the second scenario, selection would have taken place on identical haplotypes that were maintained through time in the two clusters. In our dataset, such signal of adaptive migration/selection on standing variation between clusters was observed for five genes, including TRAP that plays a key role in sporozoite motility and invasion. Although haplotypic diversity is very important in Africa, only few dominant haplotypes remain in South America, thus concomitantly confirming the strong positive selection that occurred on this gene in the New World and also suggesting that only few related haplotypes spread throughout South America through migration. It is not known how this variant was selected in this new environment. On the other hand, in Africa and Asia, TRAP is more under balancing selection whereby it maintains high levels of genetic and haplotypic diversity (Naung et al. 2022). Studies using model-based statistical approaches (Lee and Coop 2017) will be needed to investigate the different modes of convergent adaption in the American populations.

### Conclusions

We explored the genomic polymorphism of *P. falciparum* populations in the Americas and different regions of the world. By analyzing *P. falciparum* nuclear genome, we could describe its population structure and refine the history of its colonization of the Americas. We confirmed the existence of at least two independent waves of introduction from Africa: one in the North and the other in the South. Unlike previous studies, we found that populations in the SAM South cluster (Brazil – French Guiana) are the results of an ancestral admixture from the first and second waves of migration. By exploring the genomes of American populations of *P. falciparum*, we also detected many genes that are evolving under positive selection in these populations. Among them, some had already been described as selected in this continent, while others are completely new. Most genes showed only signals of selection in one cluster, suggesting that selective pressures vary among locations or that selection has not taken the same path to adapt to similar environments. However, some genes were under selection in both clusters, indicating that adaptive evolution was repeatable. For few of these genes, we found evidence that this adaptive repeated evolution occurred through adaptive migration between clusters or selection on standing ancestral variations.

## Materials and methods

### Data mapping, SNP calling, and compilation

The sequencing data for the isolates added to the *MalariaGen* dataset and the samples from the outgroup were retrieved as a FASTQ file (*Supplementary Table 1*). Then, sequencing reads were trimmed to remove adapters and preprocessed to eliminate low-quality reads (--quality-cutoff=30) using the cutadapt program (Martin 2011). Reads shorter than 50 bp and containing “N” were discarded (--minimum-length=50 --max-n=0). Sequenced reads were aligned to the Pf3D7 v3 reference genome of *P. falciparum* (Gardner et al. 2002) using bwa-mem (Li and Durbin 2009). A first filter was applied to exclude isolates with a mean genome coverage depth lower than 5×. The Genome Analysis Toolkit (GATK, version 3.8.0, McKenna et al. (2010)) was used to call SNPs in each isolate following the GATK best practices. Duplicate reads were marked using the MarkDuplicates tool from the Picard tools 2.5.0 (broadinstitute.github.io/picard/) with default options. Local realignment around indels was performed using the IndelRealigner tool from GATK. Variants were called using the HaplotypeCaller module in GATK and reads mapped with a “reads minimum mapping quality” of 30 (-mmq 30) and minimum base quality of >20 (--min_base_quality_score 20). During SNP calling, the genotypic information was kept for all sites (variants and invariant sites, --option ERC) to retain the information carried by the SNPs fixed for the reference allele. Therefore, the VCF files obtained with the *MalariaGen Project* dataset could be merged without losing information for the sites fixed in American populations and with the same nucleotide as the reference genome. VCF files were merged with BCFtools v1.10.2 (Li 2011; Danecek et al. 2021). All these steps are summarized in *Supplementary Figure S2.*

### SNP data filtration

The variant filtration steps were performed using VCFtools v 0.1.16 (Danecek et al. 2011) and BCFtools v1.10.2 (Li 2011; Danecek et al. 2021). The within-host infection complexity was assessed by calculating the *F*_WS_ values (Amegashie et al. 2020) with *vcfdo* (github.com/IDEELResearch/vcfdo; last accessed July 2022). An *F*_WS_ threshold of >0.95 was used as a proxy for monoclonal infection.

Highly related samples and clones could have generated spurious signals of population structure, biased estimators of population genetic variation, and violated the assumptions of the model-based population genetic approaches used in this study (e.g., ADMIXTURE, TreeMix, ADMIXTOOLS2) (Wang 2018). Therefore, the relatedness between haploid genotype pairs was measured by estimating the pairwise fraction of the genome identical by descent (IBD) between strains within populations using the hmmIBD program, with the default parameters for recombination and genotyping error rates, and using the allele frequencies estimated by the program (Schaffner et al. 2018). Isolate pairs that shared >50% of IBD were considered highly related. In each family of related samples, only the strain with the lowest amount of missing data was retained. All the data filtration steps are summarized in *Supplementary Figure S2.*

### Population structure, admixture, and relationships between populations

As PCA and ADMIXTURE analyses require a dataset with unlinked variants, SNPs were LD-pruned with PLINK v.2 (Chang et al. 2015). All SNPs with a correlation coefficient >0.1 (parameters: *--indep-pairwise 50 10 0.1*) were removed using a window size of 50 SNPs and a step of 10. PCA was carried out with PLINK v.2 (Chang et al. 2015) and the following parameters: *--geno --maf 0.001 --mind.* The MAF was set at 0.1% to remove doubletons. Then, the maximum likelihood clustering method implemented in the ADMIXTURE v 1.3.0 software (Alexander et al. 2009) was used with different cluster (K) numbers, from 2 to 15, with 15 replicates for each K to check consistency among replicates. The optimal K value was estimated using the cross-validation index, and convergence was checked with *pong* (Behr et al. 2016).

TreeMix and ADMIXTOOLS2 were used to estimate the most likely population tree or network topology and reticulations among them, based on variance-covariance in allele frequency. When adding the outgroup with three *P. praefalciparum* samples from Otto et al. (2018), only bi-allelic SNPs with at least no missing data in one *P. praefalciparum* sample were kept. As TreeMix and ADMIXTOOLS2 require unlinked SNPs, the dataset was LD-pruned as done for the PCA and ADMIXTURE analyses. For TreeMix, the number of migration events (*m*) that best fitted the data was calculated by running TreeMix 15 times for each *m* value, with *m* ranging from 0 to 15. The optimal *m* value (*m*=3) was estimated using the OptM R package (Fitak 2021). Then, a consensus tree with bootstrap node support was obtained by running TreeMix 100 times and post-processing using the BITE R package (Milanesi et al. 2017). To find the best network topology with ADMIXTOOLS2, the function *find_graphs* was used for 0 to 12 admixture events with 100 replicates each, and at most 300 generations for each. For the five best network topologies (*i.e.* those displaying the likelihood score closest to zero), the goodness of fit was computed with the R package *admixture-graph* (Leppälä et al. 2017). This approach allows comparing the observed *f*_4_ statistics among the different alternatives and identifying the graph(s) that best fit the data. Two graphs with the best goodness of fit and the least *f*_4_ statistics outliers were selected (*Supplementary Fig. S7*). Figure 2B shows only the graph with the lowest number of *f*_4_ statistics outliers.

### Demographic history of the American populations

The Tajima’s *D* (Tajima 1989) was measured for both American clusters, three African populations (Senegal, Democratic Republic of Congo, and Tanzania), and one Asian population (Myanmar). These populations were chosen because they had the smallest amount of missing data for each region of interest (West Africa, Central Africa, East Africa, and Asia, respectively). Tajima’s *D* values were estimated using VCFtools v 0.1.16 (Danecek et al. 2011), with a window of 5kb. The sample size was standardized (*i.e.* 20 randomly chosen isolates for each population) to obtain values that could be compared. Moreover, the variation in effective population size over time was estimated using *Stairway Plot* v2.1.1 (Liu and Fu 2020) and the same populations and clusters used to calculate the Tajima’s *D*, but without any sample size standardization. Three *P. praefalciparum* genomes from the study by Otto et al. (2018) were used to polarize the ancestral *vs.* derived states of SNPs and create an unfolded SFS. Only bi-allelic SNPs without missing data in at least one *P. praefalciparum* sample were considered for this analysis. The SFS was generated with easySFS (github.com/isaacovercast/easySFS). For these analyses, a mutation rate of 4.055×10^-9^ (Otto et al. 2018) was assumed and an observed number of sites equal to the number of *P. praefalciparum* sites that are in the core genome of this *P. falciparum* dataset (10,007,378 sites) (Miles et al. 2015).

### Detection of positive selection

Given the low sample size in each American locality (from n=5 to n=18), selection scan analyses were performed at the scale of the genetic clusters defined by ADMIXTURE and PCA, as described by Hupalo et al. (2016). Indeed, the power of *XP-EHH* and *Rsb* analyses is influenced by the sample size, and at least 20 haplotypes are recommended by Pickrell et al. (2009). Thus, each American cluster (SAM North and SAM South) was compared to the West African population from Senegal that had the smallest amount of missing data. For *ABS,* the Senegal population was kept as an outgroup and Myanmar, the Asian population with the smallest amount of missing data, was added. The *XP-EHH* and *Rsb* scores were calculated using the R package *rehh* (Gautier et al. 2017). Following Klassmann and Gautier (2022), to keep the maximum SNP number without compromising the test statistical power, data were not polarized with *P. praefalciparum*. Thus, the allele present in the reference genome was considered the ancestral allele. The significance threshold was set at -log(*p*-value) = 4, as recommended by Gautier et al. (2017), and only SNPs with negative standardized values (*i.e.* indicating positive selection for the American populations rather than African population) were considered.

The *ABS* values were calculated with *CalcABS* (Cheng et al. 2017) in sliding windows of the genome (a window of 20 kb with a step of 1 kb). All windows with <20 SNPs were removed to avoid extreme values caused by a low SNP number in some windows. The 1% most extreme values were considered as evidence that the genomic regions displayed signs of an ancient selective sweep in our South American populations. Among these extreme values, peaks (*i.e.* ≥3 consecutive windows with outliers values) were observed in some regions. Due to the large window size and to avoid high artefactual values caused by a very extreme region, only the regions with the maximum values for each peak were kept. In the absence of peaks, all points with the most extreme 1% values were kept.

Once selection signals were detected, the identified genes were annotated using the general feature format (GFF) file available from genedb (genedb.org, January 2021 version) and the intersect function of BEDtools v 2.26.0 (Quinlan and Hall 2010). Additional information (e.g. gene name, function, and biological process) was retrieved from *PlasmoDB* (plasmodb.org, accessed in February 2022).

### Detection of introgression

*RND* values (Feder et al. 2005) were calculated between SAM South and SAM North clusters, with the Senegalese population as outgroup, on sliding windows of 5kb with a step size of 1kb and at least 20 SNPs per window. To confirm the information given by the *RNDs,* the haplotype network for the three populations with extremely low *RNDs* was visualized with *popart* v1.7 (Leigh and Bryant 2015) and the Median-joining method (Bandelt et al. 1999), with ε equals zero.

## Acknowledgments

We would like to thank the *i-trop* bioinformatics platform at IRD Montpellier, the South Green Platform, and the French Bioinformatics Institute (IFB) for providing access to high-performance computing cluster (HPC) resources. We also thank the Center for Information Technology of the University of Groningen for their support and for providing access to the Peregrine high-performance computing cluster. This work was supported by the French ANR MICETRAL (ANR-19-CE35-0010) and ANR GENAD (ANR-20-CE35-003).

## Author Contributions

Conception: F.P., V.R., E.L., M.C.F. and M.J.M.L.; funding acquisition: V.R. and F.P.; method development and data analysis: M.J.M.L., J.D., M.C.F., F.P.; interpretation of the results: M.J.M.L., J.D., M.C.F., F.P., and V.R.; drafting of the manuscript: M.L.M.L. and F.P.; reviewing and editing of the manuscript: M.J.M.L., E.L., F.R., M.C.F., F.P., and V.R.

## Data Availability

The majority of data is from the *P. falciparum* Community Project conducted by MalariaGen (Pearson et al. 2019), and can be downloaded from the Wellcome Trust Sanger Institute public ftp site (ftp://ngs.sanger.ac.uk/production/malaria/pfcommunityproject/CatalogueOfVariations_v4.0/). For the other samples from Brazil, French Guiana, and Haiti, raw sequencing reads are available from the NCBI Sequencing Read Archive under the BioProject accession numbers PRJNA312679, PRJNA242163, and PRJNA603776. *P. praefalciparum* samples are accessible from the European Nucleotide Archive under sample accessions SAMEA2464702, SAMEA2073285, and SAMEA2493921. The data used in this study are available from the IRD DataSud repository (doi: *to be added*). The scripts are available in this github repository: MargauxLefebvre/P.falciparum_americas.

