## Supplementary Tables and Figures for "Population genomic evidence of adaptive response during the invasion history of *Plasmodium falciparum* in the Americas"

**Author's affiliations:**

[1] MIVEGEC, Univ. Montpellier, CNRS, IRD, Montpellier, France

[2] Malaria Biology and Vaccine Unit, Institut Pasteur, Paris, France.

[3] Groningen Institute for Evolutionary Life Sciences (GELIFES), University of Groningen, Groningen, The Netherlands

[4] REHABS, International Research Laboratory, CNRS-NMU-UCBL, George Campus, Nelson Mandela University, George, South Africa.

[§] **Co-supervised the work**

**Supplementary Table S1:** Number of *P. falciparum* isolates by country in the final dataset.

| Region | Country | Number of isolates | Sources |
|---|---|---|---|
| Central Africa | Democratic Republic of Congo | 138 | MalariaGen Project (Pearson *et al.*, 2019) |
| West Africa | Cameroon | 108 | |
| | Burkina Faso | 16 | |
| | Benin | 25 | |
| | Ghana | 364 | |
| | Guinea | 67 | |
| | Ivory Coast | 42 | |
| | Mali | 224 | |
| | Senegal | 55 | |
| | Gambia | 107 | |
| | Mauritania | 38 | |
| East Africa | Malawi | 96 | |
| | Kenya | 54 | |
| | Ethiopia | 13 | |
| | Madagascar | 19 | |
| | Tanzania | 161 | |
| | Uganda | 5 | |
| South Asia | Bangladesh | 36 | |
| West South-East Asia | Thailand | 405 | |
| | Myanmar | 103 | |
| East South-East Asia | Cambodia | 228 | |
| | Laos | 66 | |
| | Vietnam | 83 | |
| Oceania | Papua New Guinea | 77 | |
| | Indonesia | 35 | |
| Central and South Americas | Colombia | 12 | |
| | Peru | 5 | |
| | Brazil | 5 | SRA database project number PRJNA312679 (Moser *et al.*, 2020) |
| | French Guiana | 18 | SRA database project number PRJNA242163 (Pelleau *et al.*, 2015) |
| | Haiti | 8 | SRA database project number PRJNA603776 (Tagliamonte *et al.*, 2020) |

**Supplementary Table S2**: List of genes in which evidence of positive selection was observed only in the SAM North cluster (Colombia - Haiti) with *XP-EHH, Rsb* and *ABS*.

| Chr[a] | Test | ID[b] | Name | Functions | Biological processes |
|---|---|---|---|---|---|
| 1 | XP-EHH, Rsb | PF3D7_0104100 | *Unknown* | *Unknown* | Modulation by symbiont of host immune response |
| 5 | XP-EHH, Rsb | PF3D7_0511300 | *Unknown* | *Unknown* | *Unknown* |
| 5 | ABS | PF3D7_0520600 | Ap4AH | Bis(5'-nucleosyl)-tetraphosphatase (asymmetrical) activity | AMP biosynthetic process; ATP biosynthetic process |
| 5 | ABS | PF3D7_0520700 | *Unknown* | RNA polymerase II complex binding | Positive regulation of transcription elongation from RNA polymerase II promoter; recruitment of 3'-end processing factors to the RNA polymerase II holoenzyme complex |
| 5 | ABS | PF3D7_0520800 | *Unknown* | *Unknown* | *Unknown* |
| 5 | ABS | PF3D7_0520900 | SAHH | Adenosylhomocysteinase activity; protein binding | S-adenosylmethionine cycle |
| 5 | ABS | PF3D7_0521000 | *Unknown* | *Unknown* | *Unknown* |
| 5 | ABS | PF3D7_0521100 | *Unknown* | *Unknown* | *Unknown* |
| 5 | ABS | PF3D7_0521200 | *Unknown* | *Unknown* | *Unknown* |
| 6 | XP-EHH, Rsb | PF3D7_0619500 | ACS12 | Medium-chain fatty acid-CoA ligase activity | Fatty acid metabolic process |
| 6 | ABS | PF3D7_0626500 | CEP135 | *Unknown* | *Unknown* |
| 6 | ABS | PF3D7_0626600 | *Unknown* | *Unknown* | *Unknown* |
| 6 | ABS | PF3D7_0626700 | *Unknown* | ATP binding; ATP hydrolysis activity | *Unknown* |
| 6 | ABS | PF3D7_0626800 | PyrK | RNA binding; pyruvate kinase activity | Glycolytic process; protein homotetramerization |
| 6 | ABS | PF3D7_0626900 | *Unknown* | Structural constituent of ribosome | *Unknown* |
| 6 | ABS | PF3D7_0627000 | *Unknown* | *Unknown* | *Unknown* |
| 6 | ABS | PF3D7_0627100 | *Unknown* | *Unknown* | *Unknown* |
| 6 | ABS | PF3D7_0627200 | *Unknown* | *Unknown* | *Unknown* |
| 6 | ABS | PF3D7_0627300 | RNF5 | Ubiquitin protein ligase activity; ubiquitin-like protein conjugating enzyme binding; zinc ion binding | ER-associated misfolded protein catabolic process; ubiquitin-dependent protein catabolic process |
| 6 | ABS | PF3D7_0627400 | TIM22 | Mitochondrion targeting sequence binding; protein transmembrane transporter activity | Protein insertion into mitochondrial inner membrane; protein targeting to mitochondria |

| | | | | | |
|---|---|---|---|---|---|
| 6 | ABS | PF3D7_0627500 | DJ1 | Peptidase activity; protein deglycase activity | Guanine deglycation, glyoxal removal; protein deglycation; protein deglycation, glyoxal removal; protein folding; thiamine biosynthetic process |
| 6 | ABS | PF3D7_0627600 | *Unknown* | *Unknown* | *Unknown* |
| 6 | ABS | PF3D7_0627700 | *Unknown* | Nuclear import signal receptor activity; nuclear localization sequence binding; protein transmembrane transporter activity | Protein import into nucleus |
| 6 | ABS | PF3D7_0627800 | ACAS | Acetate-CoA ligase activity | Acetyl-CoA biosynthetic process; chromatin organization; generation of precursor metabolites and energy; histone acetylation; response to xenobiotic stimuli |
| 6 | ABS | PF3D7_0627900 | POP4 | Ribonuclease P RNA binding; ribonuclease activity | rRNA processing |
| 9 | ABS | PF3D7_0931900 | AKLP2 | *Unknown* | *Unknown* |
| 9 | ABS | PF3D7_0932000 | *Unknown* | *Unknown* | *Unknown* |
| 9 | ABS | PF3D7_0932100 | *Unknown* | *Unknown* | *Unknown* |
| 9 | ABS | PF3D7_0932200 | PFN | Actin binding; actin monomer binding; phospholipid binding; proline-rich region binding; protein binding | Actin cytoskeleton organization; cell motility; cytoplasmic actin-based contraction involved in cell motility; entry into host; sequestering of actin monomers |
| 9 | ABS | PF3D7_0932300 | M18AAP | *Unknown* | *Unknown* |
| 9 | ABS | PF3D7_0932400 | RF1 | *Unknown* | *Unknown* |
| 9 | ABS | PF3D7_0932500 | DHHC6 | Palmitoyltransferase activity; protein-cysteine S-palmitoyltransferase activity | Peptidyl-L-cysteine S-palmitoylation; protein palmitoylation; protein targeting to membrane |
| 9 | ABS | PF3D7_0932600 | RPS6 | Small ribosomal subunit rRNA binding; structural constituent of ribosome | *Unknown* |
| 9 | ABS | PF3D7_0932700 | *Unknown* | *Unknown* | *Unknown* |
| 9 | ABS | PF3D7_0932800 | CSE1 | Nuclear export signal receptor activity | Protein export from nucleus; protein import into nucleus |
| 9 | ABS | PF3D7_0932900 | *Unknown* | *Unknown* | *Unknown* |
| 9 | ABS | PF3D7_0933000 | *Unknown* | mRNA binding | Pre-mRNA cleavage required for polyadenylation |
| 9 | ABS | PF3D7_0933100 | *Unknown* | *Unknown* | *Unknown* |
| 9 | ABS | PF3D7_0933200 | *Unknown* | *Unknown* | Protein stabilization; regulation of protein stability |

| | | | | | |
|---|---|---|---|---|---|
| 9 | ABS | PF3D7_0933300 | *Unknown* | *Unknown* | *Unknown* |
| 9 | ABS | PF3D7_0933400 | *Unknown* | *Unknown* | *Unknown* |
| 9 | ABS | PF3D7_0933500 | *Unknown* | Gamma-tubulin binding | Cytoplasmic microtubule organization; meiotic cell cycle; microtubule nucleation by interphase microtubule organizing center; mitotic cell cycle; spindle assembly |
| 12 | XP-EHH | PF3D7_1208200 | CRMP3 | *Unknown* | Intracellular receptor signaling pathway; intracellular transport |
| 12 | Rsb | PF3D7_1239800 | *Unknown* | *Unknown* | *Unknown* |
| 13 | ABS | PF3D7_1346600 | C2AP4 | *Unknown* | *Unknown* |
| **13** | **ABS** | **PF3D7_1346700** | **P48/45** | **Protein binding** | ***Unknown*** |
| **13** | **ABS** | **PF3D7_1346800** | **P47** | **Host cell surface receptor binding** | **Evasion of host immune response; modulation by symbiont of host cellular process; suppression by symbiont of host innate immune response** |
| 13 | ABS | PF3D7_1346900 | *Unknown* | *Unknown* | *Unknown* |
| 13 | ABS | PF3D7_1347000 | WDR92 | Methylated histone binding; ubiquitin binding | Histone lysine methylation |
| 13 | ABS | PF3D7_1347100 | TOP3 | DNA binding; DNA topoisomerase activity; protein binding | DNA topological change; DNA unwinding involved in DNA replication |
| 13 | ABS | PF3D7_1347200 | NT1 | Nucleoside transmembrane transporter activity; purine nucleoside transmembrane transporter activity | Adenosine transport; purine nucleobase transport; purine nucleoside transmembrane transport; response to xenobiotic stimulus |
| 13 | ABS | PF3D7_1347300 | *Unknown* | *Unknown* | *Unknown* |
| 13 | ABS | PF3D7_1347400 | *Unknown* | *Unknown* | *Unknown* |
| 13 | ABS | PF3D7_1347500 | ALBA4 | DNA binding; RNA binding; mRNA binding; protein-containing complex binding | Regulation of translation |
| 13 | ABS | PF3D7_1347600 | *Unknown* | *Unknown* | *Unknown* |
| 13 | ABS | PF3D7_1347700 | ECT | Ethanolamine-phosphate cytidylyltransferase activity | Phospholipid biosynthetic process |
| 13 | ABS | PF3D7_1347800 | CEP72 | *Unknown* | *Unknown* |
| 13 | ABS | PF3D7_1347900 | *Unknown* | *Unknown* | *Unknown* |
| 13 | ABS | PF3D7_1348000 | *Unknown* | *Unknown* | *Unknown* |
| 13 | ABS | PF3D7_1348100 | *Unknown* | GTP binding; GTPase activity | Small GTPase mediated signal transduction |

| | | | | | |
|---|---|---|---|---|---|
| 13 | ABS | PF3D7_1348200 | *Unknown* | *Unknown* | RNA splicing |
| 13 | ABS | PF3D7_1348300 | *Unknown* | GTP binding; GTPase activity; ribosome binding; translation elongation factor activity | Mature ribosome assembly; translational elongation |
| 13 | ABS | PF3D7_1348400 | *Unknown* | *Unknown* | *Unknown* |
| 13 | ABS | PF3D7_1348500 | *Unknown* | GTPase activator activity | Activation of GTPase activity; intracellular protein transport |
| 13 | ABS | PF3D7_1348600 | *Unknown* | *Unknown* | *Unknown* |
| 13 | ABS | PF3D7_1348700 | WDR16 | *Unknown* | *Unknown* |
| 13 | ABS | PF3D7_1348800 | *Unknown* | *Unknown* | *Unknown* |
| 13 | XP-EHH, Rsb | PF3D7_1352900 | *Unknown* | *Unknown* | *Unknown* |
| 14 | ABS | PF3D7_1407600 | *Unknown* | *Unknown* | *Unknown* |
| 14 | ABS | PF3D7_1407700 | *Unknown* | *Unknown* | *Unknown* |
| 14 | ABS | PF3D7_1407800 | PM4 | RNA binding; aspartic-type endopeptidase activity | Hemoglobin catabolic process |
| 14 | ABS | PF3D7_1407900 | PMI | Aspartic-type endopeptidase activity | Hemoglobin catabolic process |
| **14** | **ABS** | **PF3D7_1408000** | **PMII** | **Aspartic-type endopeptidase activity** | **Hemoglobin catabolic process; response to xenobiotic stimuli** |
| **14** | **ABS** | **PF3D7_1408100** | **PMIII** | **Aspartic-type endopeptidase activity** | **Response to xenobiotic stimuli** |
| 14 | ABS | PF3D7_1408200 | AP2-G2 | DNA-binding transcription factor activity; protein binding; sequence-specific DNA binding | Regulation of transcription, DNA-templated |
| 14 | ABS | PF3D7_1408300 | *Unknown* | *Unknown* | *Unknown* |
| 14 | ABS | PF3D7_1408400 | FANCJ | ATP binding; DNA helicase activity; DNA polymerase binding | DNA duplex unwinding; negative regulation of DNA recombination; negative regulation of t-circle formation; regulation of double-strand break repair via homologous recombination; telomeric loop disassembly |
| 14 | ABS | PF3D7_1408500 | *Unknown* | *Unknown* | *Unknown* |
| 14 | ABS | PF3D7_1408600 | *Unknown* | RNA binding; structural constituent of ribosome | Maturation of SSU-rRNA from tricistronic rRNA transcripts (SSU-rRNA, 5.8S rRNA, LSU-rRNA); translation |
| 14 | ABS | PF3D7_1408700 | *Unknown* | K63-linked polyubiquitin modification-dependent protein binding; thiol-dependent deubiquitinase | Plastid organization; protein deubiquitination involved in ubiquitin-dependent protein catabolic processes |

[a] Chr: chromosome.
[b] Gene identifier from *Plasmodb*.
Genes discussed in this article are in bold.

**Supplementary Table S3:** List of genes where evidence of positive selection was observed only in the SAM South cluster (Brazil – French Guiana) with *XP-EHH, Rsb* and *ABS*.

| Chr[a] | Test | ID[b] | Name | Functions | Biological processes |
|---|---|---|---|---|---|
| 4 | ABS | PF3D7_0416300 | MCM9 | ATP-dependent activity, acting on DNA; DNA replication origin binding; single-stranded DNA binding | DNA replication initiation; double-strand break repair via homologous recombination |
| 4 | ABS | PF3D7_0416400 | HAT1 | H4 histone acetyltransferase activity | Histone H4 acetylation |
| 4 | ABS | PF3D7_0416500 | MAF1 | RNA polymerase III core binding | Negative regulation of transcription by RNA polymerase III |
| 4 | ABS | PF3D7_0416600 | PHBL | *Unknown* | Regulation of the mitochondrial membrane potential |
| 4 | ABS | PF3D7_0416700 | *Unknown* | *Unknown* | *Unknown* |
| 4 | ABS | PF3D7_0416800 | SAR1 | GTPase activity | Endoplasmic reticulum to Golgi vesicle-mediated transport; intracellular protein transport; membrane organization; positive regulation of protein exit from endoplasmic reticulum; regulation of COPII vesicle coating; vesicle organization |
| 4 | ABS | PF3D7_0416900 | *Unknown* | *Unknown* | *Unknown* |
| 4 | ABS | PF3D7_0417000 | *Unknown* | *Unknown* | *Unknown* |
| 4 | ABS | PF3D7_0417100 | PUF2 | RNA binding; mRNA binding; protein binding | Negative regulation of translation; posttranscriptional regulation of gene expression |
| **4** | **ABS** | **PF3D7_0417200** | **DHFR-TS** | **RNA binding; thymidylate synthase activity** | **dTMP biosynthetic process; response to drugs; response to xenobiotic stimuli** |
| 4 | ABS | PF3D7_0417300 | *Unknown* | *Unknown* | Cell metal ion homeostasis |
| 4 | ABS | PF3D7_0417400 | *Unknown* | *Unknown* | *Unknown* |
| 5 | ABS | PF3D7_0522400 | *Unknown* | *Unknown* | *Unknown* |
| 5 | ABS | PF3D7_0522500 | RPL17 | Structural constituent of ribosome | Translation |
| 5 | ABS | PF3D7_0522600 | *Unknown* | *Unknown* | *Unknown* |
| 5 | ABS | PF3D7_0522700 | SufA | 2 iron, 2 sulfur cluster binding | Iron-sulfur cluster assembly; protein maturation by iron-sulfur cluster transfer |

| | | | | | |
|---|---|---|---|---|---|
| 5 | ABS | PF3D7_0522800 | BUD31 | *Unknown* | RNA splicing; alternative mRNA splicing, via spliceosome; mRNA splicing, via spliceosome |
| 5 | ABS | PF3D7_0522900 | *Unknown* | *Unknown* | *Unknown* |
| **5** | **ABS** | **PF3D7_0523000** | **MDR1** | **ATPase-coupled transmembrane transporter activity; protein binding** | **Response to drugs; response to xenobiotic stimuli; transmembrane transport** |
| 5 | ABS | PF3D7_0523100 | QCR2 | Metalloendopeptidase activity; ubiquinol-cytochrome-c reductase activity | Protein processing involved in protein targeting to mitochondria |
| 5 | ABS | PF3D7_0523200 | *Unknown* | *Unknown* | *Unknown* |
| 5 | ABS | PF3D7_0523300 | ApiCOX18 | *Unknown* | *Unknown* |
| 5 | ABS | PF3D7_0523400 | *Unknown* | *Unknown* | *Unknown* |
| 5 | ABS | PF3D7_0523500 | *Unknown* | *Unknown* | *Unknown* |
| 5 | ABS | PF3D7_0523600 | *Unknown* | *Unknown* | *Unknown* |
| 5 | ABS | PF3D7_0523700 | *Unknown* | *Unknown* | *Unknown* |
| 7 | ABS | PF3D7_0707200 | *Unknown* | Lysine-acetylated histone binding | *Unknown* |
| 7 | ABS | PF3D7_0707300 | RAMA | Protein binding | Cell-cell adhesion; cellular protein localization; entry into host |
| 7 | ABS | PF3D7_0707400 | *Unknown* | Zinc ion binding | Mitochondrion organization |
| 7 | ABS | PF3D7_0707500 | *Unknown* | *Unknown* | *Unknown* |
| 7 | ABS | PF3D7_0707600 | MED10 | *Unknown* | *Unknown* |
| 7 | ABS | PF3D7_0707700 | *Unknown* | Chaperone binding; ubiquitin protein ligase activity | Cell response to misfolded proteins; positive regulation of proteolysis; proteasome-mediated ubiquitin-dependent protein catabolic process; protein polyubiquitination; protein quality control for misfolded or incompletely synthesized proteins |
| 7 | ABS | PF3D7_0707800 | *Unknown* | *Unknown* | *Unknown* |
| 7 | ABS | PF3D7_0707900 | *Unknown* | *Unknown* | Maturation of 5.8S rRNA; maturation of LSU-rRNA |
| 7 | ABS | PF3D7_0708000 | *Unknown* | Alpha-tubulin binding | Microtubule cytoskeleton organization; post-chaperonin tubulin folding pathway; tubulin complex assembly |

| | | | | | |
|---|---|---|---|---|---|
| 7 | ABS | PF3D7_0708100 | RPB10 | DNA-directed 5'-3' RNA polymerase activity; zinc ion binding | tRNA transcription by RNA polymerase III; transcription by RNA polymerase I; transcription by RNA polymerase II |
| 7 | ABS | PF3D7_0708200 | *Unknown* | *Unknown* | *Unknown* |
| 7 | ABS | PF3D7_0708300 | BUD32 | Protein binding; protein serine/threonine kinase activity | Protein phosphorylation; tRNA threonylcarbamoyladenosine metabolic process |
| 7 | ABS | PF3D7_0708400 | HSP90 | ATP hydrolysis activity; RNA binding; protein binding; unfolded protein binding | Cell response to heat; protein folding; protein stabilization |
| 7 | ABS | PF3D7_0708500 | *Unknown* | *Unknown* | Regulation of gene expression; response to heat; response to unfolded protein |
| 7 | ABS | PF3D7_0708600 | IMC1d | *Unknown* | *Unknown* |
| 7 | ABS | PF3D7_0708700 | COX4 | *Unknown* | *Unknown* |
| 7 | ABS | PF3D7_0708800 | HSP110c | ATP hydrolysis activity; protein binding | Protein folding; response to heat |
| 7 | ABS | PF3D7_0708900 | SCO1 | *Unknown* | Mitochondrial cytochrome C oxidase assembly |
| **7** | **ABS** | **PF3D7_0709000** | **CRT** | **Ferric iron transmembrane transporter activity; xenobiotic transmembrane transporter activity** | **Glutathione transport; peptide transport; response to drugs; response to xenobiotic stimuli** |
| 7 | ABS | PF3D7_0709100 | *Unknown* | *Unknown* | *Unknown* |
| 7 | ABS | PF3D7_0709200 | GLP3 | *Unknown* | *Unknown* |
| 7 | ABS | PF3D7_0709300 | MED14 | *Unknown* | *Unknown* |
| 7 | ABS | PF3D7_0709400 | *Unknown* | *Unknown* | *Unknown* |
| 7 | ABS | PF3D7_0709500 | *Unknown* | *Unknown* | *Unknown* |
| 7 | ABS | PF3D7_0709700 | PARE | Acylglycerol lipase activity; hydrolase activity; hydrolase activity, acting on ester bonds; lipase activity | Monoacylglycerol catabolic process; phospholipid metabolic process; response to xenobiotic stimuli |
| 7 | ABS | PF3D7_0709800 | *Unknown* | *Unknown* | *Unknown* |
| 7 | XP-EHH, Rsb | PF3D7_0710100 | *Unknown* | *Unknown* | *Unknown* |
| 8 | XP-EHH, Rsb | PF3D7_0809200 | pfa55-14 | Histone acetyltransferase activity; peptide alpha-N-acetyltransferase activity | N-terminal peptidyl-methionine acetylation; histone H3 acetylation; histone H4 acetylation |
| 8 | XP-EHH, Rsb | PF3D7_0809400 | *Unknown* | *Unknown* | *Unknown* |

| | | | | | |
|---|---|---|---|---|---|
| 8 | ABS | PF3D7_0815200 | *Unknown* | Nuclear import signal receptor activity; nuclear localization sequence binding | NLS-bearing protein import into nucleus; protein import into nucleus |
| 8 | ABS | PF3D7_0815300 | *Unknown* | *Unknown* | *Unknown* |
| 8 | ABS | PF3D7_0815400 | ATPTG3 | *Unknown* | *Unknown* |
| 8 | ABS | PF3D7_0815500 | *Unknown* | *Unknown* | *Unknown* |
| 8 | ABS | PF3D7_0815600 | EIF3G | mRNA binding; translation initiation factor activity | Translation; translational initiation |
| 8 | ABS | PF3D7_0815700 | Ub | ATP hydrolysis activity | Regulation of cell cycle process; ubiquitin-dependent ERAD pathway |
| 8 | ABS | PF3D7_0815800 | VPS9 | Guanyl-nucleotide exchange factor activity | Golgi to endosome transport |
| 8 | ABS | PF3D7_0815900 | aLipDH | Dihydrolipoyl dehydrogenase activity; flavin adenine dinucleotide binding; protein homodimerization activity | Acetyl-CoA biosynthetic process from pyruvate; oxidation-reduction process |
| 8 | ABS | PF3D7_0816000 | RRB1 | *Unknown* | Ribosome biogenesis |
| 8 | ABS | PF3D7_0816100 | PPCDC | FMN binding; phosphopantothenoylcysteine decarboxylase activity | Coenzyme A biosynthetic process |
| 8 | ABS | PF3D7_0816200 | VPS2 | *Unknown* | Endosome transport via multivesicular body sorting pathway; late endosome to vacuole transport; protein transport |
| 8 | ABS | PF3D7_0816300 | *Unknown* | *Unknown* | *Unknown* |
| 8 | ABS | PF3D7_0816400 | *Unknown* | Calcium ion binding | *Unknown* |
| 8 | ABS | PF3D7_0816500 | HSP20 | Protein self-association; unfolded protein binding | Protein complex oligomerization; protein folding; response to heat; response to hydrogen peroxide; response to reactive oxygen species; response to salt stress; response to unfolded proteins |
| 8 | ABS | PF3D7_0816600 | ClpB1 | ATP hydrolysis activity | Cellular response to heat; response to unfolded proteins |
| 8 | ABS | PF3D7_0816700 | TRAPPC2L | *Unknown* | Endoplasmic reticulum to Golgi vesicle-mediated transport |
| 8 | ABS | PF3D7_0816800 | DMC1 | ATP-dependent activity, acting on DNA; DNA strand exchange activity; double-stranded DNA binding; single-stranded DNA binding | DNA recombinase assembly; chromosome organization involved in meiotic cell cycle; mitotic recombination; reciprocal meiotic recombination; response to ionizing radiation; strand invasion |

| | | | | | |
|---|---|---|---|---|---|
| 8 | ABS | PF3D7_0816900 | AK2 | Adenylate kinase activity; protein binding | Nucleoside diphosphate metabolic process; purine nucleotide metabolic process |
| 8 | ABS | PF3D7_0817000 | UBA3 | NEDD8 activating enzyme activity | Protein modification by small protein conjugation; protein neddylation |
| 8 | ABS | PF3D7_0817100 | *Unknown* | GTPase activity; ferrous iron transmembrane transporter activity | tRNA wobble uridine modification |
| 8 | ABS | PF3D7_0817200 | *Unknown* | *Unknown* | *Unknown* |
| 8 | ABS | PF3D7_0817300 | *Unknown* | *Unknown* | *Unknown* |
| 8 | ABS | PF3D7_0817400 | *Unknown* | Double-stranded DNA binding; polydeoxyribonucleotide 5'-hydroxyl-kinase activity; polynucleotide 3'-phosphatase activity | DNA repair; nucleotide phosphorylation |
| 8 | ABS | PF3D7_0823400 | *Unknown* | *Unknown* | *Unknown* |
| 8 | ABS | PF3D7_0823500 | IMC1i | *Unknown* | *Unknown* |
| 8 | ABS | PF3D7_0823600 | LipB | Lipoate-protein ligase activity | Lipoate biosynthetic process; protein lipoylation |
| 8 | ABS | PF3D7_0823700 | TOM7 | *Unknown* | Protein targeting to mitochondria |
| 8 | ABS | PF3D7_0823800 | *Unknown* | RNA binding; unfolded protein binding | Chaperone cofactor-dependent protein refolding; protein refolding |
| 8 | ABS | PF3D7_0823900 | DTC | Dicarboxylic acid transmembrane transporter activity; oxoglutarate:malate antiporter activity | Alpha-ketoglutarate transport; dicarboxylic acid transport; malate transport; mitochondrial transport |
| 8 | ABS | PF3D7_0824000 | *Unknown* | *Unknown* | *Unknown* |
| 8 | ABS | PF3D7_0824100 | *Unknown* | *Unknown* | *Unknown* |
| 8 | ABS | PF3D7_0824200 | *Unknown* | *Unknown* | *Unknown* |
| 8 | ABS | PF3D7_0824300 | Obg2 | GTP binding | *Unknown* |
| 8 | ABS | PF3D7_0824400 | NT2 | Nucleoside transmembrane transporter activity | Nucleoside transport |
| 8 | ABS | PF3D7_0824500 | *Unknown* | *Unknown* | *Unknown* |
| 8 | ABS | PF3D7_0824600 | DRE2 | *Unknown* | Iron-sulfur cluster assembly |
| 8 | ABS | PF3D7_0824700 | LMF1 | *Unknown* | *Unknown* |
| 8 | ABS | PF3D7_0824800 | *Unknown* | *Unknown* | *Unknown* |
| 8 | ABS | PF3D7_0824900 | *Unknown* | *Unknown* | *Unknown* |

| | | | | | |
|---|---|---|---|---|---|
| 8 | ABS | PF3D7_0825000 | *Unknown* | *Unknown* | *Unknown* |
| 8 | ABS | PF3D7_0825100 | *Unknown* | *Unknown* | *Unknown* |
| 8 | ABS | PF3D7_0825200 | IF3a | Ribosome binding; translation initiation factor activity | Ribosome disassembly |
| 8 | ABS | PF3D7_0825300 | *Unknown* | *Unknown* | *Unknown* |
| 10 | ABS | PF3D7_1036500 | *Unknown* | *Unknown* | *Unknown* |
| 10 | ABS | PF3D7_1036600 | *Unknown* | *Unknown* | *Unknown* |
| 10 | ABS | PF3D7_1036700 | PhLP2 | *Unknown* | *Unknown* |
| 10 | ABS | PF3D7_1036800 | ACT | Solute:proton symporter activity | Response to xenobiotic stimuli |
| 10 | ABS | PF3D7_1037100 | PyKII | Pyruvate kinase activity | Glycolytic process |
| 10 | ABS | PF3D7_1037200 | *Unknown* | *Unknown* | *Unknown* |
| 10 | ABS | PF3D7_1037300 | AAC1 | ATP:ADP antiporter activity; RNA binding | *Unknown* |
| 10 | ABS | PF3D7_1037400 | *Unknown* | Ubiquitin protein ligase activity | Protein monoubiquitination; protein polyubiquitination |
| 10 | ABS | PF3D7_1037500 | DYN2 | GTPase activity; microtubule binding | Mitochondrial fission; organelle fission |
| 10 | ABS | PF3D7_1037600 | XPB | 3'-5' DNA helicase activity; ATP hydrolysis activity; DNA binding; DNA helicase activity; damaged DNA binding; helicase activity; transcription factor binding | Nucleotide-excision repair, DNA duplex unwinding; nucleotide-excision repair, DNA incision; response to UV; transcription by RNA polymerase II; transcription initiation from RNA polymerase II promoter |
| 10 | ABS | PF3D7_1037700 | ERH | mRNA binding | *Unknown* |
| 10 | ABS | PF3D7_1037800 | *Unknown* | Dynein heavy chain binding; dynein light chain binding | Cilium movement; microtubule-based movement |
| 12 | ABS | PF3D7_1225100 | api-IRS | Isoleucine-tRNA ligase activity | Isoleucyl-tRNA aminoacylation; response to xenobiotic stimuli |
| 12 | ABS | PF3D7_1225200 | *Unknown* | *Unknown* | *Unknown* |
| 12 | ABS | PF3D7_1225300 | *Unknown* | Small ribosomal subunit rRNA binding | *Unknown* |
| 12 | ABS | PF3D7_1225400 | *Unknown* | *Unknown* | *Unknown* |
| 12 | ABS | PF3D7_1225500 | *Unknown* | rRNA (pseudouridine) methyltransferase activity; rRNA binding | rRNA base methylation |
| 12 | ABS | PF3D7_1225600 | *Unknown* | *Unknown* | *Unknown* |
| 12 | ABS | PF3D7_1225700 | *Unknown* | *Unknown* | Phosphatidylinositol biosynthetic process; positive regulation of kinase activity |

| | | | | | |
|---|---|---|---|---|---|
| 12 | ABS | PF3D7_1225800 | UBA1 | Ubiquitin activating enzyme activity | Cell response to DNA damage stimulus; protein modification by small protein conjugation; protein ubiquitination; ubiquitin-dependent protein catabolic process |
| 12 | ABS | PF3D7_1225900 | *Unknown* | *Unknown* | *Unknown* |
| 12 | ABS | PF3D7_1226000 | *Unknown* | *Unknown* | *Unknown* |
| 12 | ABS | PF3D7_1226100 | HAD3 | Catalytic activity; hydrolase activity; magnesium ion binding; phosphatase activity | *Unknown* |
| 12 | ABS | PF3D7_1226200 | *Unknown* | *Unknown* | *Unknown* |
| 12 | ABS | PF3D7_1226300 | HAD2 | *Unknown* | *Unknown* |
| 12 | ABS | PF3D7_1226400 | *Unknown* | *Unknown* | Regulation of autophagy |
| 12 | ABS | PF3D7_1243400 | *Unknown* | *Unknown* | *Unknown* |
| 12 | ABS | PF3D7_1243500 | SNF7 | *Unknown* | Late endosome to vacuole transport via multivesicular body sorting pathway; vesicle budding from membrane |
| 12 | ABS | PF3D7_1243600 | *Unknown* | RNA binding; translation initiation factor activity | Translational initiation |
| 12 | ABS | PF3D7_1243700 | *Unknown* | Ubiquitin conjugating enzyme activity | Cell response to DNA damage stimulus; protein polyubiquitination |
| 12 | ABS | PF3D7_1243800 | WDR82 | Chromatin binding | Histone H3-K4 methylation; histone H3-K4 trimethylation |
| 12 | ABS | PF3D7_1243900 | DOC2 | Calcium-dependent phospholipid binding | Entry into host |
| 12 | ABS | PF3D7_1244000 | *Unknown* | Flavin adenine dinucleotide binding | Mitochondrial tRNA wobble uridine modification; tRNA methylation; tRNA wobble uridine modification |
| 12 | ABS | PF3D7_1244100 | *Unknown* | *Unknown* | N-terminal peptidyl-methionine acetylation |
| 12 | ABS | PF3D7_1244200 | TFB2 | Double-stranded DNA binding | Nucleotide-excision repair; phosphorylation of RNA polymerase II C-terminal domain; transcription by RNA polymerase II |
| 12 | ABS | PF3D7_1244400 | *Unknown* | *Unknown* | *Unknown* |
| 12 | ABS | PF3D7_1244500 | PIMMS57 | *Unknown* | *Unknown* |
| 12 | ABS | PF3D7_1244600 | ARFGAP1 | DNA binding; GTPase activator activity | COPI coating of Golgi vesicles |
| 14 | ABS | PF3D7_1440800 | MFS6 | *Unknown* | *Unknown* |

| | | | | | |
|---|---|---|---|---|---|
| 14 | ABS | PF3D7_1440900 | *Unknown* | *Unknown* | *Unknown* |
| 14 | ABS | PF3D7_1441000 | NCS2 | Sulfurtransferase activity | tRNA thio-modification; tRNA wobble position uridine thiolation |
| 14 | ABS | PF3D7_1441100 | *Unknown* | *Unknown* | *Unknown* |
| 14 | ABS | PF3D7_1441200 | *Unknown* | RNA binding; structural constituent of ribosome | Maturation of LSU-rRNA; translation |
| 14 | ABS | PF3D7_1441300 | *Unknown* | Protein serine/threonine kinase activity | Protein phosphorylation |
| 14 | ABS | PF3D7_1441400 | FACT-S | DNA binding; RNA binding; histone binding; nucleosome binding; single-stranded DNA binding | *Unknown* |
| 14 | ABS | PF3D7_1441500 | *Unknown* | *Unknown* | *Unknown* |
| 14 | ABS | PF3D7_1441600 | *Unknown* | *Unknown* | *Unknown* |
| 14 | ABS | PF3D7_1441700 | ATP23 | *Unknown* | Mitochondrial protein processing; mitochondrial proton-transporting ATP synthase complex assembly |
| 14 | ABS | PF3D7_1441800 | VPS60 | *Unknown* | Late endosome to vacuole transport via the multivesicular body sorting pathway; vesicle budding from membrane |
| 14 | ABS | PF3D7_1441900 | TFB5 | *Unknown* | Nucleotide-excision repair; nucleotide-excision repair, preincision complex assembly; phosphorylation of RNA polymerase II C-terminal domain; transcription by RNA polymerase II |
| 14 | ABS | PF3D7_1442000 | *Unknown* | GTP binding; GTPase activity | Positive regulation of microtubule polymerization; protein ADP-ribosylation |
| 14 | ABS | PF3D7_1442100 | RPA3 | Damaged DNA binding; single-stranded DNA binding | DNA replication; base-excision repair; double-strand break repair via homologous recombination; mismatch repair; nucleotide-excision repair |
| 14 | ABS | PF3D7_1442200 | YihA2 | GTPase activity; mitochondrial ribosome binding | *Unknown* |
| 14 | ABS | PF3D7_1442300 | tRIP | RNA binding; aminoacyl-tRNA ligase activity | tRNA aminoacylation for protein translation |
| 10 | ABS | PF3D7_1036900 | *Unknown* | mRNA binding | *Unknown* |
| 10 | ABS | PF3D7_1037000 | REV3 | DNA-directed DNA polymerase activity | Double-strand break repair via homologous recombination; error-prone translesion synthesis |

[a] Chr: chromosome.
[b] Gene identifier In *Plasmodb*.
Genes in bold are discussed in this article.

**Supplementary Table S4**: List of genes where evidence of positive selection was detected in both South American clusters with *XP-EHH, Rsb* and *ABS*.

| Chr[a] | North[b] | South[c] | ID[d] | Name | Functions | Processes |
|---|---|---|---|---|---|---|
| 5 | ABS | ABS | PF3D7_0521300 | *Unknown* | *Unknown* | *Unknown* |
| 5 | ABS | ABS | PF3D7_0521400 | *Unknown* | *Unknown* | *Unknown* |
| 5 | ABS | ABS | PF3D7_0521500 | *Unknown* | Pseudouridine synthase activity; pseudouridylate synthase activity | Enzyme-directed rRNA pseudouridine synthesis; rRNA processing |
| 5 | ABS | ABS | PF3D7_0521600 | *Unknown* | *Unknown* | *Unknown* |
| 5 | ABS | ABS | PF3D7_0521700 | DDX1 | *Unknown* | rRNA processing |
| 5 | ABS | ABS | PF3D7_0521800 | *Unknown* | ATP hydrolysis activity | *Unknown* |
| 5 | ABS | ABS | PF3D7_0521900 | *Unknown* | *Unknown* | *Unknown* |
| 5 | ABS | ABS | PF3D7_0522000 | *Unknown* | *Unknown* | *Unknown* |
| 5 | ABS | ABS | PF3D7_0522100 | *Unknown* | *Unknown* | *Unknown* |
| 5 | ABS | ABS | PF3D7_0522200 | TAF10 | Protein binding | Transcription by RNA polymerase II |
| 5 | ABS | ABS | PF3D7_0522300 | *Unknown* | S-adenosylmethionine-dependent methyltransferase activity; rRNA (guanine) methyltransferase activity | rRNA (guanine-N7)-methylation |
| 6 | XP-EHH, Rsb | XP-EHH, Rsb | PF3D7_0619600 | *Unknown* | *Unknown* | *Unknown* |
| 7 | XP-EHH, Rsb | XP-EHH, Rsb | PF3D7_0704600 | UT | Ubiquitin-protein transferase activity | Response to drugs; response to xenobiotic stimuli |
| 7 | Rsb | ABS | PF3D7_0709600 | POP1 | Ribonuclease MRP activity; ribonuclease P activity | RNA phosphodiester bond hydrolysis, endonucleolytic; tRNA processing |
| 7 | XP-EHH | ABS | PF3D7_0709900 | *Unknown* | *Unknown* | *Unknown* |
| 7 | XP-EHH, Rsb | XP-EHH, RSb | PF3D7_0710000 | *Unknown* | rRNA binding | Ribosomal large subunit assembly |

| | | | | | | |
|---|---|---|---|---|---|---|
| 7 | XP-EHH, Rsb | XP-EHH, Rsb | PF3D7_0710200 | *Unknown* | *Unknown* | *Unknown* |
| 7 | XP-EHH, Rsb | XP-EHH, Rsb | PF3D7_0710400 | RAD14 | Damaged DNA binding | UV-damage excision repair; base-excision repair; nucleotide-excision repair involved in interstrand cross-link repair; nucleotide-excision repair, DNA damage recognition; nucleotide-excision repair, DNA incision |
| 7 | XP-EHH, Rsb | XP-EHH, Rsb | PF3D7_0711000 | CDC48 | ATP hydrolysis activity; polyubiquitin modification-dependent protein binding; ubiquitin binding | ER-associated misfolded protein catabolic process; autophagosome maturation; macroautophagy; mitotic spindle disassembly; regulation of cell cycle process; retrograde protein transport, ER to cytosol; ubiquitin-dependent ERAD pathway |
| 7 | XP-EHH, Rsb | XP-EHH, Rsb | PF3D7_0711200 | *Unknown* | *Unknown* | *Unknown* |
| 7 | XP-EHH, Rsb | XP-EHH | PF3D7_0711500 | *Unknown* | Chromatin binding | Ran protein signal transduction; chromosome condensation; chromosome segregation; regulation of mitotic spindle assembly |
| 7 | XP-EHH, Rsb | XP-EHH, Rsb | PF3D7_0713500 | *Unknown* | *Unknown* | *Unknown* |
| 7 | XP-EHH, Rsb | XP-EHH, Rsb | PF3D7_0713600 | *Unknown* | Structural constituent of ribosome | Translation |
| 7 | XP-EHH, Rsb | XP-EHH, Rsb | PF3D7_0713900 | *Unknown* | N-acetyltransferase activity | *Unknown* |
| 8 | XP-EHH, Rsb | XP-EHH, Rsb | PF3D7_0809600 | *Unknown* | Cysteine-type endopeptidase activity | Meiotic chromosome separation |
| 8 | ABS | ABS | PF3D7_0817700 | RON5 | RNA binding; host cell surface binding | *Unknown* |
| 8 | ABS | ABS | PF3D7_0817800 | C3AP3 | *Unknown* | *Unknown* |

| | | | | | | |
|---|---|---|---|---|---|---|
| 8 | ABS | ABS | PF3D7_0817900 | HMGB2 | DNA binding; DNA binding, bending | Positive regulation of cytokine production involved in immune response; regulation of transcription by RNA polymerase II; regulation of transcription by RNA polymerase III; regulation of transcription, DNA-templated |
| 8 | ABS | ABS | PF3D7_0818000 | SNRNP27 | *Unknown* | *Unknown* |
| 8 | ABS | ABS | PF3D7_0818100 | *Unknown* | *Unknown* | *Unknown* |
| 8 | ABS | ABS | PF3D7_0818200 | 14-3-3I | RNA binding; histone binding; protein binding | Entry into host |
| 8 | ABS | ABS | PF3D7_0818300 | *Unknown* | Dynein complex binding | Mitotic spindle organization |
| 8 | ABS | ABS | PF3D7_0818400 | FCF1 | *Unknown* | *Unknown* |
| 8 | ABS | ABS | PF3D7_0818500 | *Unknown* | Zinc ion binding | *Unknown* |
| 8 | ABS | ABS | PF3D7_0818600 | PBLP | Lipase activity; palmitoyl-(protein) hydrolase activity | Cell maturation; entry into host; lipid metabolic process |
| 8 | ABS | ABS | PF3D7_0818700 | *Unknown* | ATP-dependent activity, acting on DNA; DNA binding; chromatin binding | ATP-dependent chromatin remodeling |
| **11** | **XP-EHH, Rsb** | **Rsb** | **PF3D7_1133400** | **AMA1** | **Host cell surface binding; protein binding** | **Entry into host** |
| 12 | ABS | ABS | PF3D7_1206800 | *Unknown* | *Unknown* | *Unknown* |
| 12 | ABS | ABS | PF3D7_1206900 | *Unknown* | *Unknown* | *Unknown* |
| 12 | ABS | ABS | PF3D7_1207000 | *Unknown* | *Unknown* | *Unknown* |
| 12 | ABS | ABS | PF3D7_1207100 | *Unknown* | RNA binding | rRNA processing |
| 12 | ABS | ABS | PF3D7_1207200 | *Unknown* | *Unknown* | *Unknown* |
| 12 | ABS | ABS | PF3D7_1207300 | LIMP | *Unknown* | Cell gliding; entry into host |
| 12 | ABS | ABS | PF3D7_1207400 | *Unknown* | *Unknown* | *Unknown* |
| 12 | ABS | ABS | PF3D7_1207500 | *Unknown* | RNA binding | *Unknown* |

| | | | | | | |
|---|---|---|---|---|---|---|
| 12 | ABS | ABS | PF3D7_1207600 | MiaA | ATP binding; tRNA binding; tRNA dimethylallyltransferase activity | tRNA modification; tRNA processing |
| 12 | ABS | ABS | PF3D7_1207700 | *Unknown* | *Unknown* | *Unknown* |
| 12 | ABS | ABS | PF3D7_1207800 | *Unknown* | *Unknown* | *Unknown* |
| 12 | ABS | ABS | PF3D7_1207900 | *Unknown* | *Unknown* | *Unknown* |
| 12 | ABS | ABS | PF3D7_1208000 | *Unknown* | *Unknown* | *Unknown* |
| **13** | **XP-EHH, Rsb** | **XP-EHH, Rsb** | **PF3D7_1335900** | **TRAP** | **Host cell surface receptor binding** | **Cell gliding; entry into host; movement of cell or subcellular component** |

[a] Chr: chromosome

[b] Test in which this signal was found in the SAM North cluster (Colombia - Haiti).

[c] Test in which this signal was found in the SAM South cluster (Brazil - French Guiana).

[d] Gene identifier in *Plasmodb*.

Genes in bold are discussed in this article.

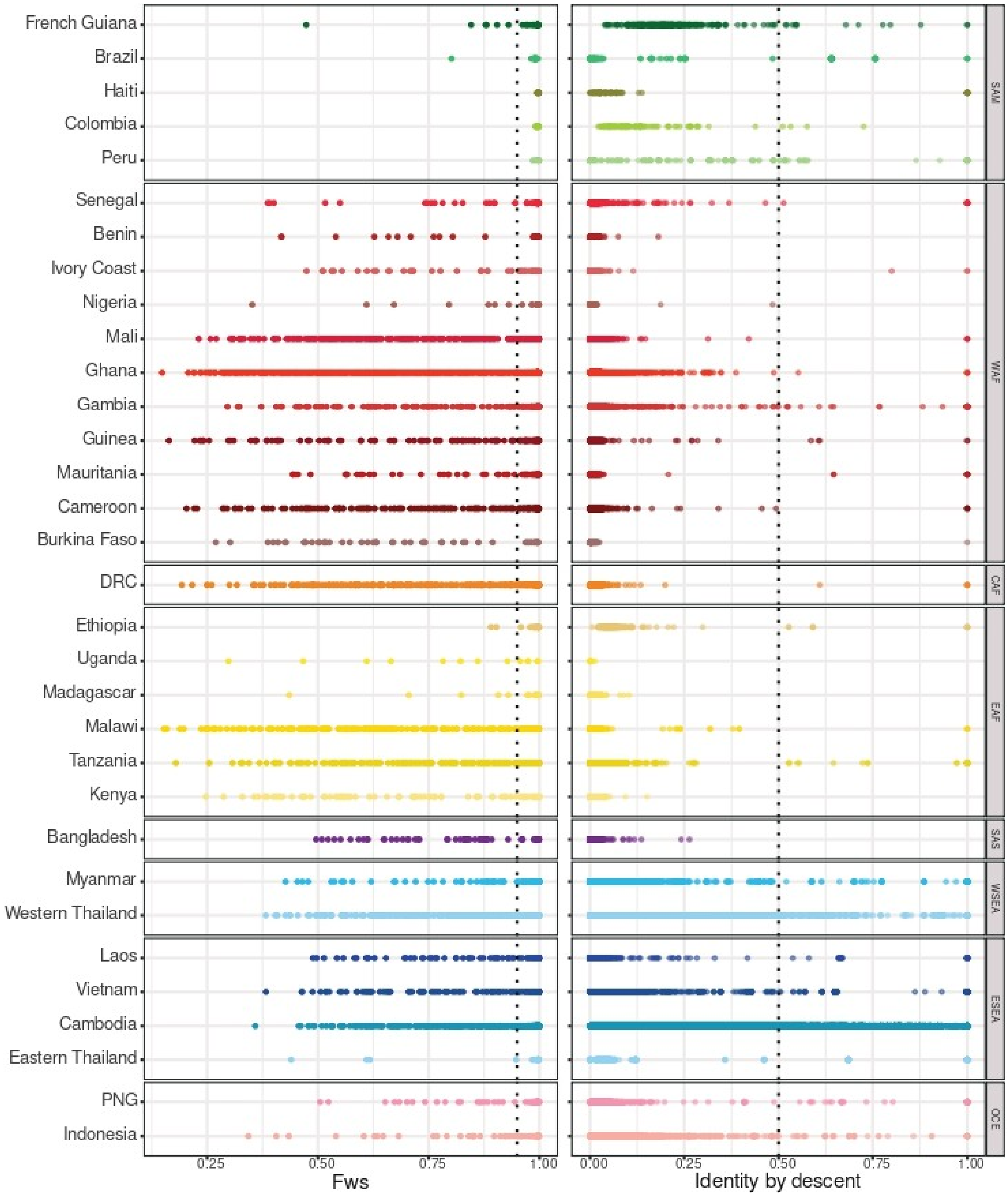


**Supplementary Figure S1: Within-sample infection complexity ($F_{WS}$ index) and consanguinity (identity by descent) in *P. falciparum* populations.** The $F_{WS}$ index provides a proxy of the diversity within individual infections, from 0 (high diversity) to 1 (no diversity). $F_{WS}$ values >0.95 (dotted line) usually indicate monoclonal infections. Identity by descent (IBD) indicates the percentage of the genome resulting from inbreeding between pairs of individuals of the same population. One individual was excluded for all strain pairs displaying a pairwise-IBD >0.5 (dotted line).

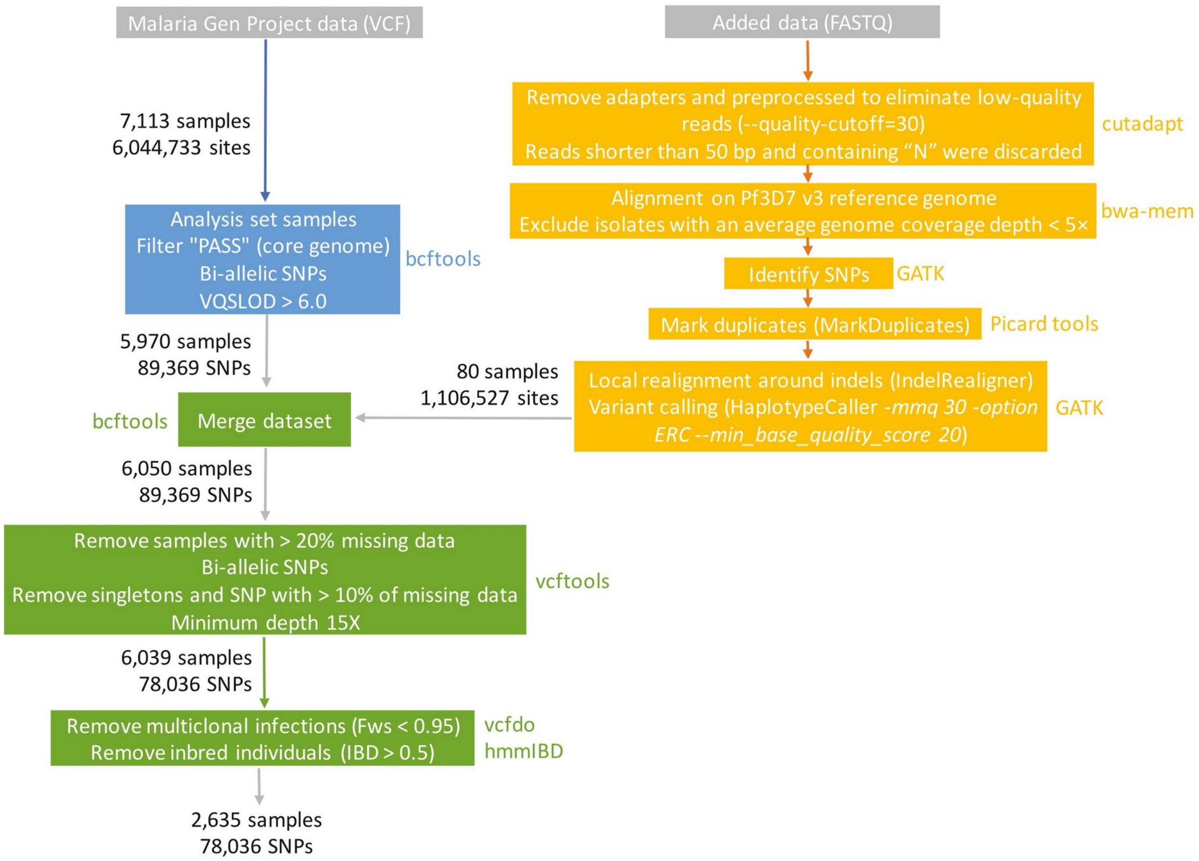


**Supplementary Figure S2: Filtering and dataset creation steps.** Each box represents a filtering step and the tool used. After each filtering step, the number of remaining samples and sites/SNPs is specified.

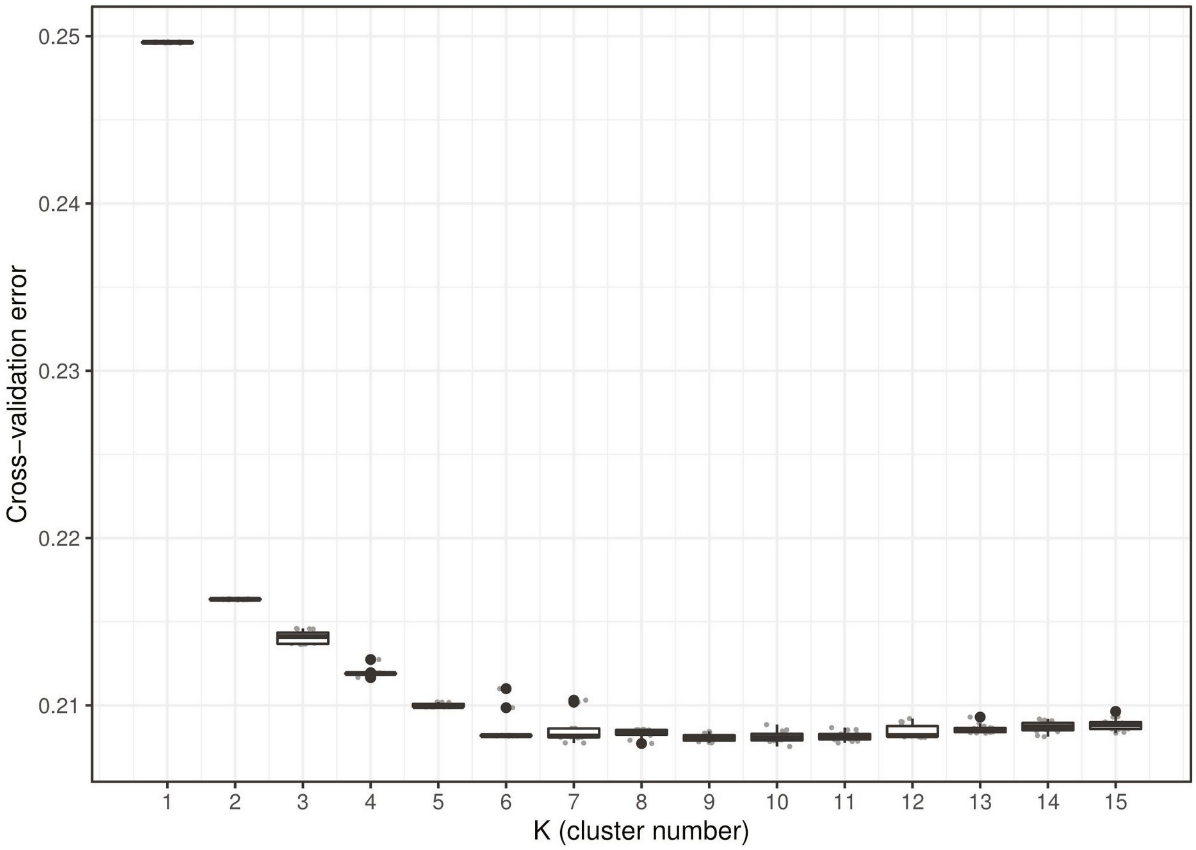


**Supplementary Figure S3: Box plot displaying the cross-validation error rate in function of the increasing number of clusters (K).** With K between 1 and 15 (15 replicates for each K value), K=9 was chosen to analyze the SNP data, as the value that minimizes the cross-validation error rate.

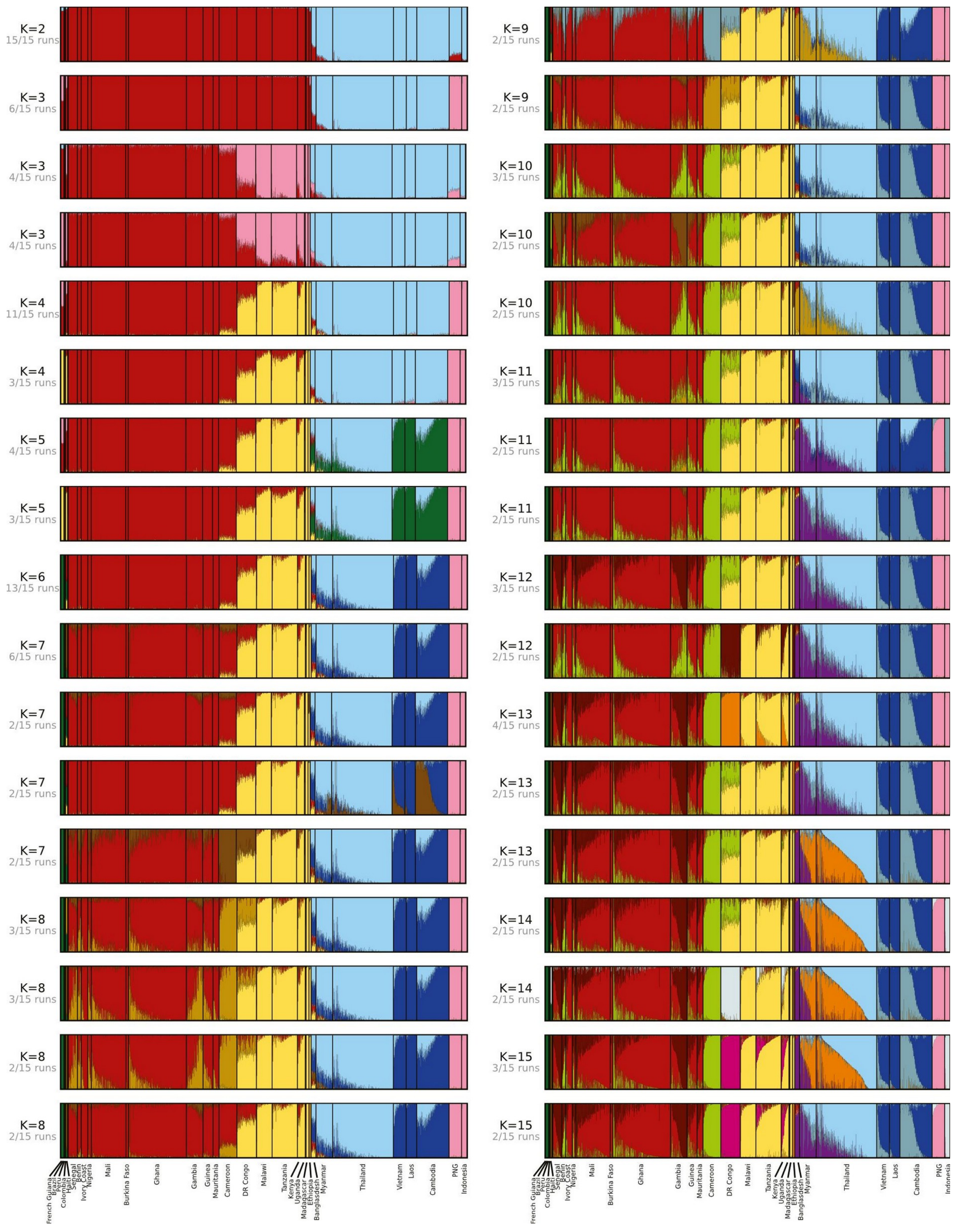


**Supplementary Figure S4: Genetic ancestry of *P. falciparum* populations worldwide estimated with ADMIXTURE (K=2 to K=15).** The number of clusters tested is described by the K value on the left, with the number of congruent runs. Only results with more than one congruent run are shown.

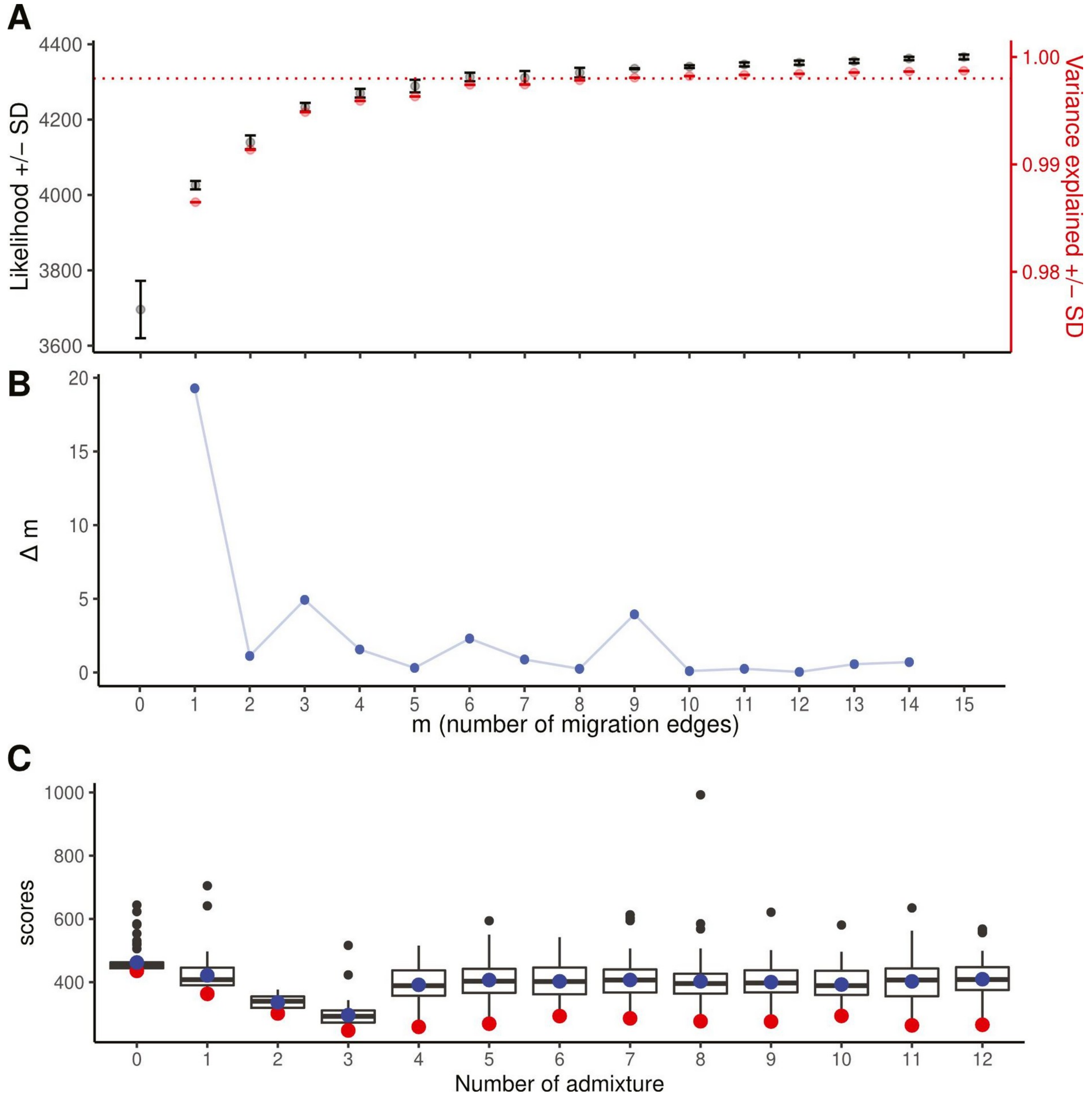


**Supplementary Figure S5: Identification of the optimal number of migration edges in *P. falciparum* populations worldwide. A.** Changes in the mean likelihood score (± SD) and the mean total fraction of the genetic variance explained (± SD) in function of the number of migration edges (TreeMix analysis). **B.** The second-order rate of change in likelihood (Δm) across values of migration edges (m). For both panels, the OptM package and the Evanno method were used, with 15 replicates for each migration edge number, from 0 to 15. The inflection point for both statistics was observed at m=3. **C.** Boxplots of the likelihood score for each number of admixture events, with 100 replicates in the ADMIXTOOLS2 analysis. Blue dots represent the mean, and red dots the minimum value.

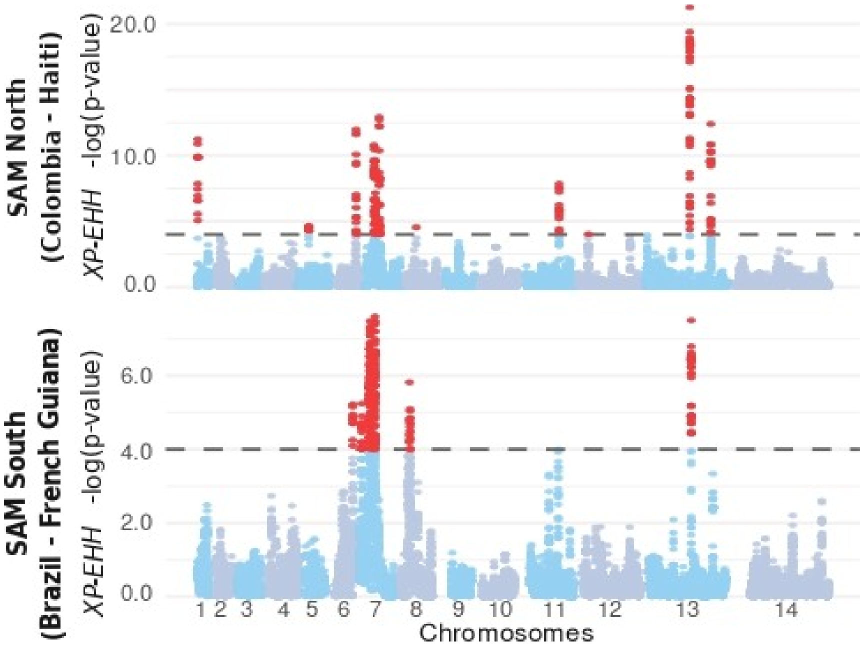

**Supplementary Figure S6:** Manhattan plots showing the *XP-EHH* scores for SAM North (Colombia - Haiti) and SAM South (Brazil - French Guiana), compared with the African reference population (Senegal). The dotted lines represent the significance threshold value of -log(p-value) = 4. Red, SNPs marking a selective sweep in the SAM cluster (negative *XP-EHH* values).

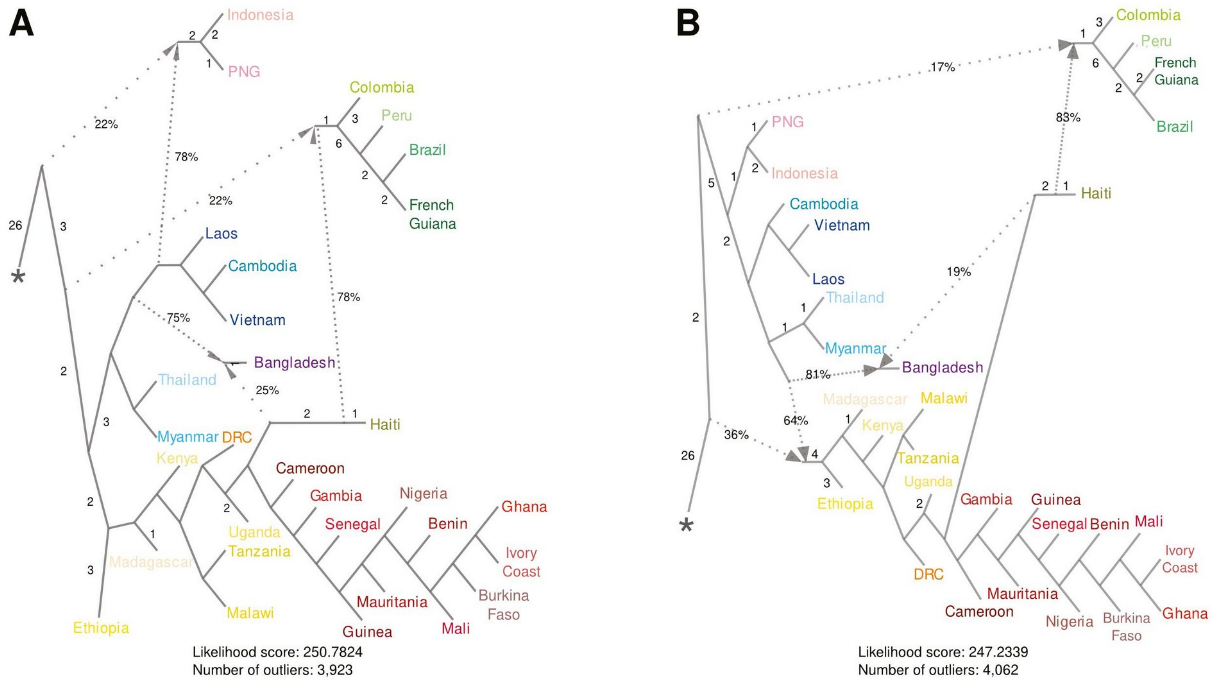


**Supplementary Figure S7: Best ADMIXTOOLS2 trees. A.** Tree with the best goodness of fit (fewest number of $f_4$ statistics outliers over 107,880 $f_4$ statistics in total) and the second best likelihood score. This tree is the one presented in Figure 2. **B.** Tree with the second best goodness of fit (second fewer number of $f_4$ statistics outliers over 107,880 $f_4$ statistics in total) and best likelihood score.
